# Cell type-specific role of CBX2-cPRC1 at the onset of spermatogonial differentiation

**DOI:** 10.1101/2022.08.25.505322

**Authors:** Jongmin J. Kim, Emma R. Steinson, Mei Sheng Lau, David C. Page, Robert E. Kingston

## Abstract

Polycomb group (PcG) proteins maintain the repressed state of lineage-inappropriate genes and are therefore essential for embryonic development and adult tissue homeostasis. One critical function of PcG complexes is modulating chromatin structure. Canonical Polycomb repressive complex 1 (cPRC1), particularly its component CBX2, can compact chromatin and phase separate *in vitro*. These activities are hypothesized to be critical for forming a repressed physical environment in cells. While much has been learned by studying these PcG activities in cell culture models, it is largely unexplored how cPRC1 regulates adult stem cells and their subsequent differentiation during tissue homeostasis in living animals. Here, we show that CBX2 is upregulated as spermatogonial stem cells differentiate and is required in the differentiating spermatogonia of the male germ line. CBX2 forms condensates, similar to previously described Polycomb bodies, that co-localize with repressed target genes in differentiating spermatogonia. Single cell analyses of mosaic *Cbx2* mutant testes show CBX2 is specifically required to produce differentiating, A1 spermatogonia. Furthermore, the domain of CBX2 responsible for compaction and phase separation is needed for the long-term maintenance of male germ cells in the animal. These results emphasize that the regulation of chromatin structure by CBX2 at a specific stage of spermatogenesis is critical for the continued production of sperm and, ultimately, the transmission of genetic material to the next generation.

## Introduction

Polycomb Group (PcG) proteins are critical for the stable inheritance of the repressed state of gene expression in development. By regulating chromatin structure and modifications of target genes, PcG proteins create a heritable repressed state over multiple divisions of cells that lack the signal that generated the repressed state (Blackledge and Klose, 2021). Rapidly proliferating cells in fly embryos can maintain their positional identity because PcG proteins prevent the misexpression of inappropriate identity genes (Struhl and Akam, 1985). The core function of PcG proteins is utilized in diverse biological processes across distant multicellular species. For example, PcG proteins are required for X chromosome inactivation in mammals (Plath et al., 2003; Silva et al., 2003; Wang et al., 2001) and for vernalization by stable silencing of a floral repressor to induce flowering (Gendall et al., 2001). This maintenance property is broadly used and is particularly important for developmental transitions, such as in stem cell self-renewal versus differentiation (Boyer et al., 2006; Endoh et al., 2008) or in cell fate bifurcations (Hirabayashi et al., 2009; Iovino et al., 2013), where one fate is stabilized over the other.

Adult stem and progenitor cells require PcG proteins both for their maintenance and differentiation (Flora et al., 2021). Many tissues in adult animals, such as blood and skin, are constantly replenished by rare adult stem cells. These cells are capable of self-renewal and can make all differentiated cell types in the lineage. Genetic ablation of certain core PcG proteins, such as *Ring1a/b* or *Eed*, results in failure of maintenance of blood (Piunti et al., 2014; Xie et al., 2014), skin (Dauber et al., 2016), intestine (Chiacchiera et al., 2016; Koppens et al., 2016), and male germ line lineages (Maezawa et al., 2017; Mu et al., 2014). Some PcG proteins repress premature senescence to facilitate adult stem cell maintenance (Lessard and Sauvageau, 2003; Molofsky et al., 2003; Park et al., 2003). As cancers often originate from misregulated adult stem and progenitor cells, mutations in PcG genes are associated with different types of human cancers (Parreno et al., 2022). The cellular mechanisms of PcG proteins’ role in tissue homeostasis and cancer are poorly understood. This is partly because the complex behavior of adult stem and progenitor cells in their native tissue environment cannot be easily modeled with the cell culture systems. Adult stem and progenitor cells are heterogeneous with different rates of proliferation and self-renewal capacity (Li and Clevers, 2010). Understanding potentially distinct functions of PcG proteins in different cellular contexts in an animal is critical for delineating the physiological effects that result from specific PcG dysfunction.

To achieve heritable gene silencing, PcG proteins covalently modify histones and modulate chromatin structure. PcG proteins form three major types of stable multi-protein complexes in the nucleus. Non-canonical Polycomb Repressive Complex 1 (ncPRC1, also called variant PRC1) catalyzes mono-ubiquitylation of Lys119 residue of H2A (H2AK119Ub1) (de Napoles et al., 2004; Wang et al., 2004), while PRC2 catalyzes mono-, di-, and tri-methylation of Lys27 residue of H3 (H3K27me1/2/3) (Fig. 1A) (Cao et al., 2002; Czermin et al., 2002; Kuzmichev et al., 2002; Muller et al., 2002). Canonical PRC1 (cPRC1), the focus of this study, is composed of RING1B (or paralog RING1A), PCGF2 (or paralog PCGF4), CBX2 (or paralogs CBX4/6/7/8), and PHC2 (or paralogs PHC1/3) (Fig. 1A) (Gao et al., 2012; Levine et al., 2002; Shao et al., 1999). cPRC1 is recruited to genes with the H3K27me3 modification catalyzed by PRC2 (Cao *et al*., 2002; Fischle et al., 2003; Min et al., 2003). cPRC1 can compact polynucleosome arrays (Francis et al., 2004; Grau et al., 2011), phase separate *in vitro* (Plys et al., 2019; Seif et al., 2020; Tatavosian et al., 2019) and mediate long-range interactions between distant PcG target loci (Isono et al., 2013; Kundu et al., 2017). These activities contribute to the formation of membraneless nuclear structures called Polycomb bodies (Saurin et al., 1998), which are proposed to be critical for the memory of gene repression (Bantignies et al., 2011; Isono *et al*., 2013). Mutations disrupting the compaction or oligomerization activities of cPRC1 result in developmental defects (Isono *et al*., 2013; Lau et al., 2017), suggesting critical non-enzymatic roles of cPRC1.

**Figure 1.**
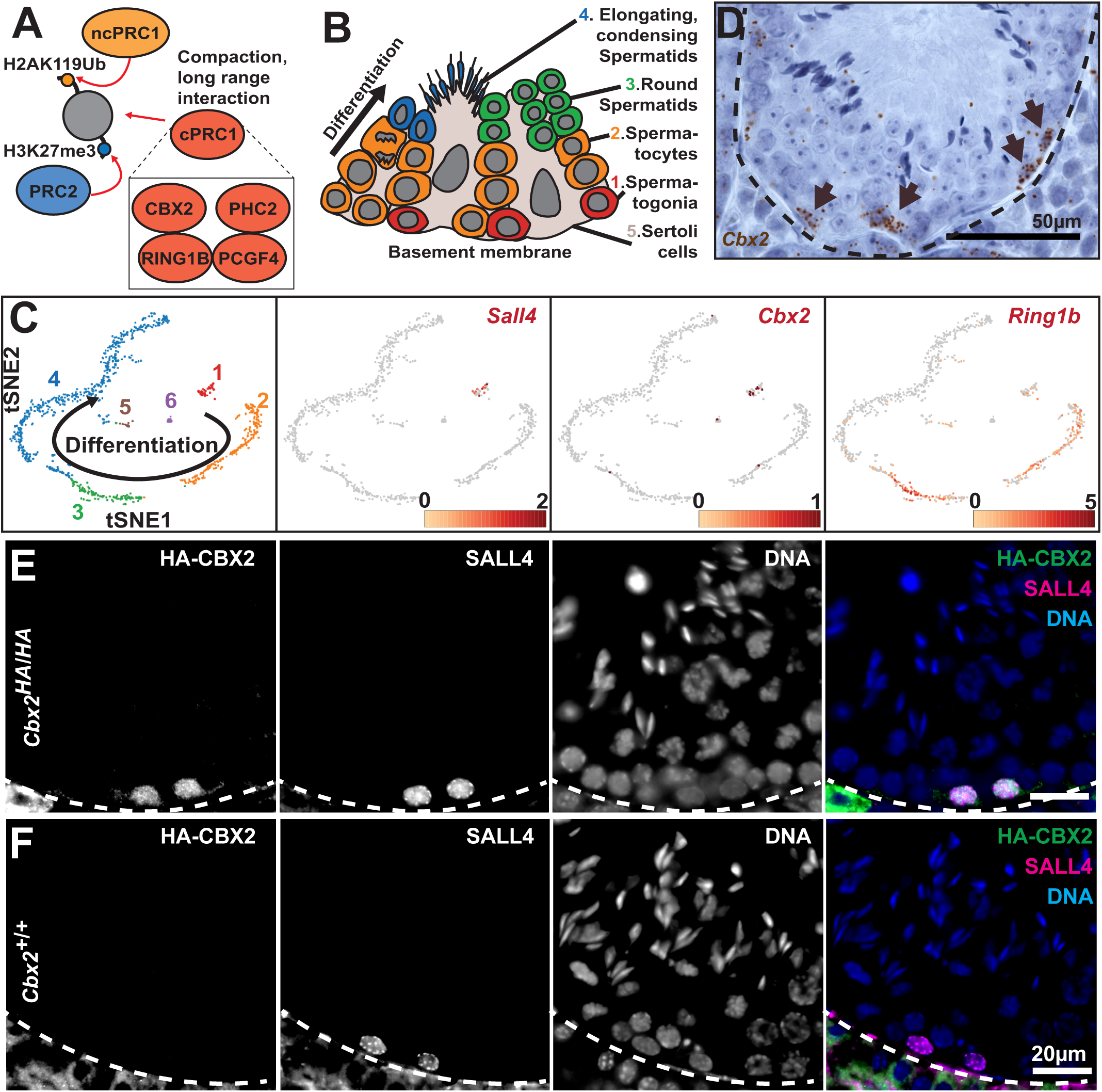
CBX2 is expressed in spermatogonia (A) Schematic representing PcG complexes’ functions on chromatin. Individual components of cPRC1 were depicted in the rectangle. (B) Schematic of cellular organization in the seminiferous tubule. Note, to show all the major steps in spermatogenesis in one picture, this schematic does not reflect stereotypical cellular organizations based on different tubule stages. (C) t-SNE representation of scRNA-seq results from adult testes. The arrow indicates the direction of germ cell differentiation based on known marker gene expression. Numbers (1-5) correspond to different cell types depicted in (B). (6) represents Leydig cells. Normalized expression levels of *Sall4*, *Cbx2* and *Ring1b* are represented in red. Color bars represent expression levels. (D) RNA *in situ* hybridization of *Cbx2* using branched signal amplification with each brown dot representing a single *Cbx2* RNA transcript. The black dotted line represents basement membrane of a seminiferous tubule. Brown arrows indicate *Cbx2*(+) cells residing at the basement membrane (E and F) Co-immunofluorescence staining of testis sections of (E) *Cbx2^HA/HA^*and (F) *Cbx2^+/+^* (negative control) animals with HA and SALL4 antibodies. Dotted lines represent the basement membrane. Images are a part of the stage XII tubules.

CBX2 is a key component of cPRC1 that is required for its non-enzymatic repressive function. cPRC1 or CBX2 by itself can compact nucleosome arrays, inhibit chromatin remodeling by mSWI/SNF complexes and phase separate together with nucleosome arrays *in vitro* (Grau *et al*., 2011; Plys *et al*., 2019; Tatavosian *et al*., 2019). These activities are dependent on a positively charged and disordered region called the Compaction and Phase Separation (CaPS, Fig. S1A) domain of CBX2 (Grau *et al*., 2011; Jaensch et al., 2021). This domain is also required for proper axial development of mice (Lau *et al*., 2017). Positive charge and disorderedness are conserved features for the compaction activity of other PcG proteins in different species (Beh et al., 2012). Thus, multivalent interactions between PcG proteins, such as CBX2, and nucleosomes through unstructured, charged residues are one important component of PcG activity. CBX2 is essential for many developmental processes. >90% of *Cbx2* mutant mice died before weaning with diverse developmental defects in axial patterning (Core et al., 1997), splenic vasculature (Katoh-Fukui et al., 2005), bone growth (Katoh-Fukui et al., 2019) and sex determination (Garcia-Moreno et al., 2019; Katoh-Fukui et al., 1998). However, in many cases, it remains an open question if and what specific cell types in development are susceptible to the loss of CBX2.

We set out to test two major questions; whether CBX2 is required at a specific stage of adult stem cell differentiation, and whether the compaction and phase separation function of CBX2 is critical for that process. Germ cells in the testis provide an ideal system to delineate the role of CBX2-cPRC1 function in adult tissue homeostasis. The male germ line is a stereotypical adult stem cell lineage, producing sperm essential for continuation of life. A small number of spermatogonial stem cells (SSCs) produce many differentiated sperm throughout the life of the organism. A key regulatory point in spermatogenesis is the irreversible commitment into ’differentiating spermatogonia’ (de Rooij and Russell, 2000). As the daughter cells of SSC divisions become ’A aligned’ spermatogonia, they gradually lose their stem cell potential. They downregulate SSC-enriched genes, such as *Foxc2* (Wei et al., 2018), and reciprocally upregulate early differentiation genes, such as *Rarg* (*retinoic acid receptor γ*) (Gely-Pernot et al., 2012; Ikami et al., 2015). When stimulated by retinoic acid (RA), some cells from this pool of SSCs and A aligned spermatogonia irreversibly differentiate to ’differentiating spermatogonia’ (Fig. 2A) (Koubova et al., 2006; Zhou et al., 2008). An important question is how a subset of these cells acquire competence to differentiate by RA stimulation, because the balance between maintenance of SSC pool and differentiation is key to continued production of sperm (Endo et al., 2015). Given the known role for PcG proteins in cell fate stabilization, we reasoned CBX2 may play a role in regulation of spermatogonial differentiation.

**Figure 2.**
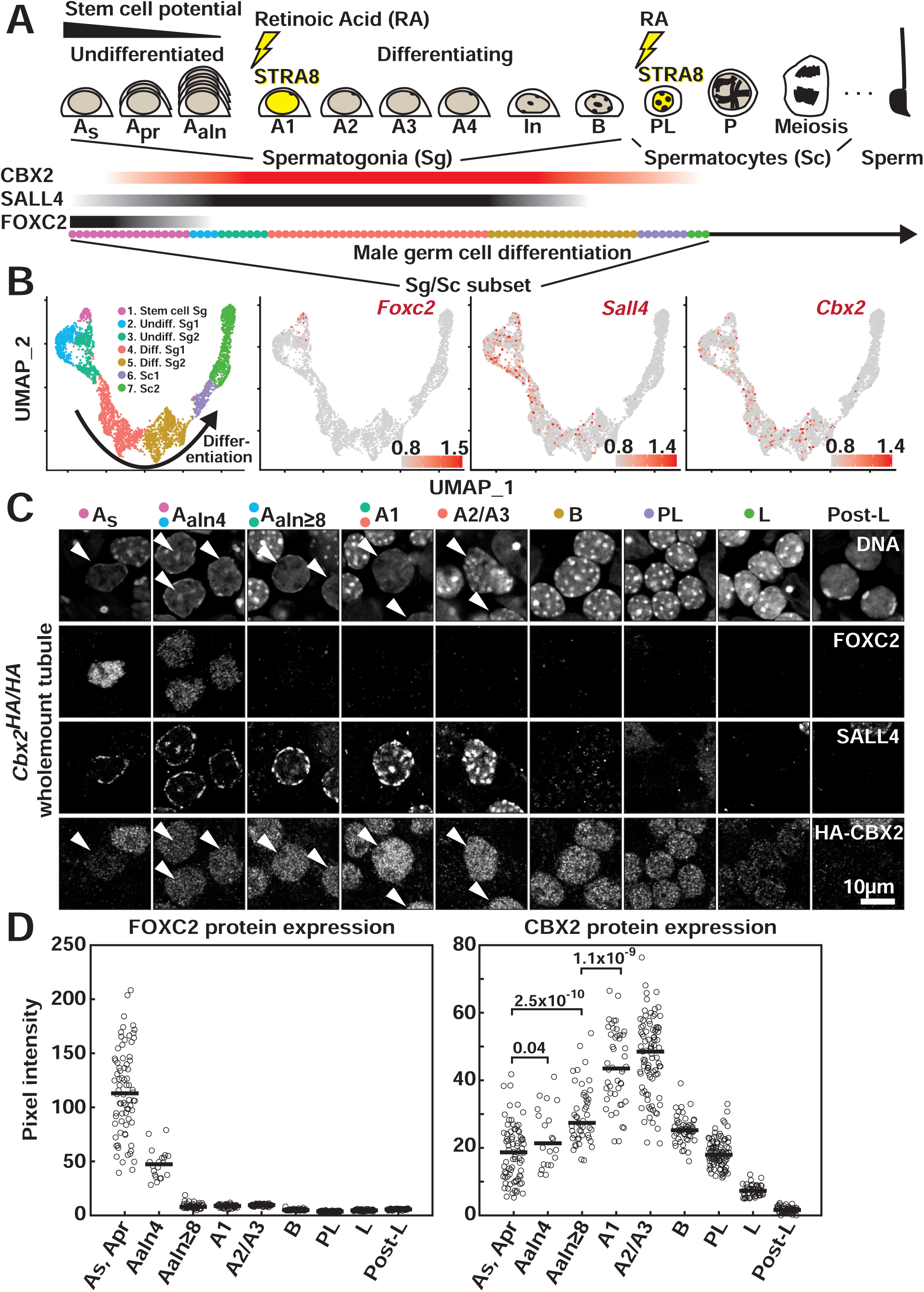
CBX2 is upregulated as spermatogonial stem cells initiate differentiation (A) Schematic of spermatogonial differentiation in mouse spermatogenesis (As: A single, Apr: A paired, Aaln: A aligned, In: Intermediate, PL: Pre Leptotene, P: Pachytene). Yellow lightning signals represent developmental transitions induced by retinoic acid and the ensuing upregulation of STRA8 (<u>St</u>imulated by <u>R</u>etinoic <u>A</u>cid 8). Cell types expressing *Foxc2*, *Sall4*, and *Cbx2* are marked by black and red bars. Colored dots at the bottom represent corresponding cell types identified in scRNA-seq analyses in (B). (B) UMAP representation of scRNA-seq results from postnatal day 15 (p15) mouse testes (Ernst *et al*., 2019). P15 UMAP was plotted after obtaining a subset of spermatogonia and early spermatocyte populations from the whole p15 dataset. P15 subset cells clustered into 7 populations and cell types were assigned according to marker gene expression (marker profiles in Fig. S2A). Normalized expression levels of *Foxc2*, *Sall4* and *Cbx2* are represented in red. (C) Co-immunofluorescence staining of whole testis tubules of *Cbx2^HA/HA^*animals with HA, SALL4, and FOXC2 antibodies. Cellular stages were identified based on the cellular organization along the tubule, the nuclear morphology based on DNA staining, and the protein expression patterns of marker genes, such as FOXC2 and SALL4. (D) Quantification of FOXC2 and CBX2 protein expression based on wholemount tubule stainings. Numbers between groups As, Apr, Aaln4, and Aaln≥8 represent *p* values calculated by two-tailed t-tests.

Here, through utilizing inducible as well as CaPS domain-specific *Cbx2* mutant mice, we show that CBX2 function is specifically required at the developmental transition from stem cells to committed progenitor cells in spermatogenesis. CBX2 expression was upregulated as germ cells initiate differentiation from spermatogonial stem cells to differentiating spermatogonia. Single cell RNA-seq combined with *Cbx2* genotyping using mosaic *Cbx2* mutant animals revealed that CBX2 function was specifically required for production of committed progenitor cells. Furthermore, the compaction and phase separation domain of CBX2 was required for germ cell maintenance. We hypothesize that CBX2 is required for stable repression of the spermatogonial stem cell program to provide competence for the cells to differentiate into committed progenitors.

## Results

### CBX2 expression coincides with the onset of spermatogonial differentiation

The testis contains spermatogonial stem cells and a continuous stream of differentiating daughter germ cells that eventually become sperm, in addition to somatic supporting cells (Fig. 1B). To identify the cell types that require CBX2 function in spermatogenesis, we profiled the expression of *Cbx2* RNA and protein in the mouse testis. Single cell RNA-seq of a wild-type testis showed *Cbx2* expression was enriched in spermatogonia (cluster #1, *Sall4*(+)), while a core PRC1 component, *Ring1b*, was broadly expressed in spermatogonia, spermatocytes and round spermatids (Fig. 1B and 1C). Consistent with the single cell RNA-seq, RNA *in situ* hybridization showed that *Cbx2* mRNA signal was detected in a small number of cells at the basement membrane, where spermatogonia reside (Fig. 1D). To detect CBX2 protein, we generated a knock-in mouse line in which two tandem HA epitopes were inserted in frame into the N-terminus of the endogenous *Cbx2* locus (*HA-Cbx2* mouse, see legends; Fig. S1A and S1B). Co-immunostaining of the spermatogonial marker SALL4 and HA-CBX2 in testis sections revealed that CBX2 was expressed in SALL4(+) spermatogonia (Fig. 1E and F). All SALL4(+) cells expressed CBX2, but CBX2 was also expressed in more differentiated pre-leptotene spermatocytes that lack SALL4 expression (Fig. S1D; stage VII-VIII, arrows), and becomes undetectable in leptotene spermatocytes (Fig. S1D; stage IX, arrowheads).

Spermatogonia can be further divided into ‘undifferentiated’ spermatogonia including spermatogonial stem cells (SSCs), and ‘differentiating’ spermatogonia committed to differentiation (Fig. 2A). As these spermatogonia have distinct properties, such as self-renewal, proliferation rate, and retinoic acid responsiveness, we set out to determine the precise stages of spermatogenesis where CBX2 is expressed. Since some PRC1 components, such as BMI1 (PCGF4), have been proposed to be a specific marker for SSCs (Komai et al., 2014), we were particularly curious if CBX2 is expressed in SSCs or not. To this end, we first used publicly available single cell RNA-seq data from juvenile mouse testes which contain a higher proportion of spermatogonia (Ernst et al., 2019). We chose cells classified as spermatogonia and early spermatocytes, and re-clustered them to generate seven refined clusters of the early cell types (Fig. 2B and S2A). Consistent with our single cell RNA-seq data from an adult testis, *Cbx2* was expressed in *Sall4*(+) spermatogonia and was downregulated at spermatocyte stages (Fig. 2B and S2B). Notably, *Cbx2* expression appeared depleted in *Foxc2*(+) spermatogonia known to have high stem cell potential (Fig. 2B) (Wei *et al*., 2018). Only 4% of *Foxc2*(+) cluster #1 cells contained any *Cbx2* reads, while *Foxc2*-low cluster #2 and #3 each had 18% and 23% of cells with *Cbx2* reads (Fig. S2B).

Coimmunostaining of CBX2 and FOXC2 also showed lower CBX2 protein expression in undifferentiated FOXC2(+) cells than differentiating spermatogonia. We performed wholemount tubule staining (Fig. S2C), because unlike in sections, FOXC2(+) cells can be categorized into singlets and paired cells (high stem cell potential) to a chain of aligned cells (low stem cell potential). Isolated and paired cells with robust FOXC2 expression showed undetectable to very low CBX2 signal (Fig. 2C and 2D; As, Apr). In contrast, aligned FOXC2(+) cells showed higher levels of CBX2 expression than the FOXC2(+) As and Apr cells (Fig. 2C and 2D; Aaln4 and Aaln≥8). CBX2 expression was further increased in differentiating spermatogonia (Fig. 2C and 2D; A1 and A2/A3), perdured until pre-leptotene spermatocytes (Fig. 2C and 2D; PL), and nearly disappeared as cells matured into the leptotene spermatocyte stage (Fig. 2C and 2D; L). We conclude that CBX2 is upregulated as spermatogonia exit from the stem cell state, which coincides with a decrease in the FOXC2 expression, and robustly expressed in differentiating spermatogonia, but is downregulated as meiosis initiates in leptotene spermatocytes.

### CBX2-cPRC1 forms Polycomb bodies in spermatogonia

Chromatin bound cPRC1 coalesces in membraneless structures called Polycomb bodies (Satijn et al., 1997; Saurin *et al*., 1998). Although it has been proposed that Polycomb bodies are critical for genome organization and PcG target gene regulation (Bantignies *et al*., 2011), there is limited evidence if Polycomb bodies are formed in primary cells of adult mammals. To ask if CBX2-cPRC1 forms Polycomb bodies in spermatogonia, we performed co-immunostaining of CBX2 and other cPRC1 proteins using dissociated germ cells. Fine sub-nuclear structures like Polycomb bodies are better visualized as a dissociated monolayer in single cells than in the intact tissue. We used FACS to purify differentiating spermatogonia with robust CBX2 expression using a cell surface marker c-KIT from *Cbx2^HA/+^* animals (c-KIT expression is specific to differentiating spermatogonia, Fig. S2A) (Shinohara et al., 2000). Co-immunostaining of the c-KIT(+) spermatogonia with HA and RING1B antibodies showed non-uniform expression of HA-CBX2 in the nucleus forming 200∼400nm (diameter) sized puncta (Fig. 3A). In addition, RING1B, which forms cPRC1 with CBX2, also showed a punctate expression pattern, overlapping with CBX2 Polycomb bodies. Not all CBX2 and RING1B puncta overlap, which is in part due to non-specific staining of antibodies especially for weak intensity puncta (Fig. S3A). To corroborate the findings using the HA tag, we developed specific polyclonal antibodies against near-full length CBX2 protein (Fig. S1A and S1C). Immunostaining of native CBX2 from wild-type c-KIT(+) spermatogonia also showed CBX2 forming Polycomb bodies together with another cPRC1 component PHC2 (Fig. 3B). Thus, CBX2-containing cPRC1 is compartmentalized into Polycomb bodies in spermatogonia.

**Figure 3.**
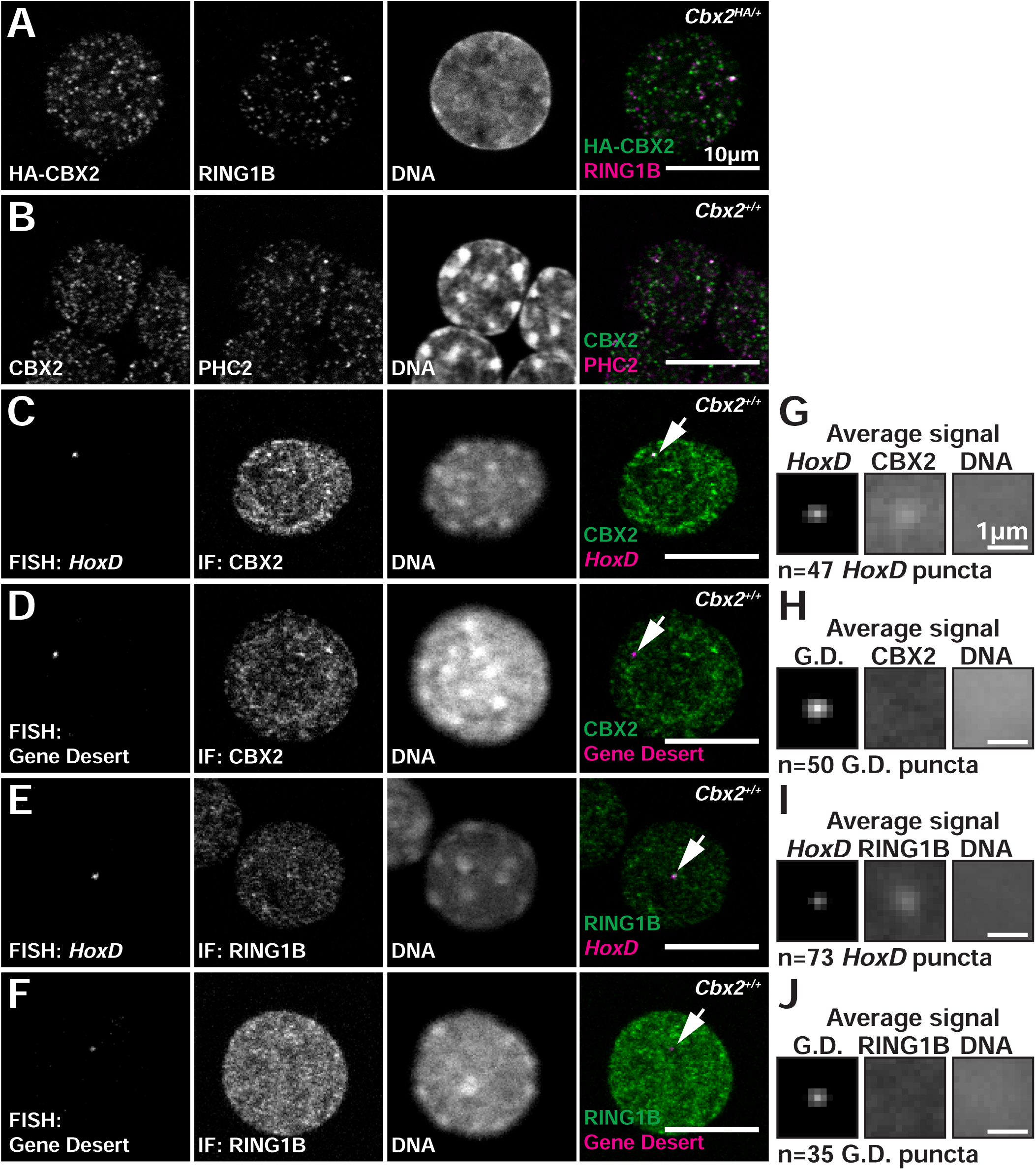
CBX2-cPRC1 forms Polycomb bodies with target DNA (A and B) Co-immunostaining of (A) HA-CBX2 and RING1B and (B) CBX2 and PHC2 using FACS sorted c-KIT(+) spermatogonia from (A) *Cbx2^HA/+^* and (B) wild-type mice. (C-F) Co-immuno-FISH of (C) *HoxD* locus and CBX2 protein, (D) gene desert locus and CBX2 protein, (E) *HoxD* locus and RING1B protein, and (F) gene desert locus and RING1B protein using FACS sorted c-KIT(+) spermatogonia from wild-type mice. A single optical section, which usually contains one allele of *HoxD* or gene desert locus in the plane, was represented to avoid incidental overlap of puncta at different focal planes. (G-J) Average signal projection of 2 µm square centered at *HoxD* or gene desert puncta. Average signal projection of CBX2 or RING1B and DNA at the corresponding locations were also represented.

One critical question is whether CBX2 Polycomb bodies represent CBX2-cPRC1 bound to chromatin, or on the contrary a storage unit sequestered away from target genes as some reports hypothesized (Saurin *et al*., 1998). Polycomb bodies were shown to be associated with target DNA, such as a *Hox* locus in fly embryos (Grimaud et al., 2006) and mouse embryonic fibroblasts (MEFs) (Isono *et al*., 2013), but whether that is true in adult animals has not been addressed. We tested whether CBX2 and RING1B condensates overlap with a classical Polycomb target gene, *HoxD*. A non-target, gene desert region was also probed as a negative control. CUT&RUN experiments using purified spermatogonia (see details in the next section) showed that CBX2 and RING1B were preferentially localized at the *HoxD* locus but not at the gene desert region (Fig. S3B). Co-immuno-FISH revealed that both CBX2 and RING1B condensates overlapped with the *HoxD* but not with the gene desert locus (Fig. 3C-F). Not all *HoxD* puncta overlapped CBX2 and RING1B condensates. About 1/3 of *HoxD* puncta overlapped with distinct CBX2 or RING1B puncta, 1/3 of *HoxD* puncta overlapped with diffusive CBX2 or RING1B signal, and the remaining 1/3 of *HoxD* puncta did not overlap with CBX2 or RING1B signal (Fig. S3C and S3D). Importantly, the negative control gene desert puncta never overlapped with CBX2 or RING1B puncta (Fig. 3D, 3F, and S3D). When images centered at *HoxD* or gene desert puncta were averaged, there was corresponding enrichment of CBX2 or RING1B signal at the *HoxD* signal peak, whereas there was no such enrichment for the gene desert locus (Fig. 3G-J). Another *Hox* Cluster, *HoxB* also showed similar CBX2 signal enrichment when *HoxB* puncta signals were averaged (Fig. S3E and S3F), while a negative control protein, SALL4, did not show signal enrichment corresponding to *HoxB* puncta locations. Thus, CBX2 not only forms Polycomb bodies with other cPRC1 components in spermatogonia, these Polycomb bodies are associated with target chromatin that harbor silenced genes.

### CBX2-cPRC1 binds genes expressed in spermatogonial stem cells and in embryonic tissues

The data presented above are consistent with CBX2-cPRC1 being necessary for repression of genes that might impede proper differentiation of spermatogonia. To examine this hypothesis, we identified genes in spermatogonia that were occupied by CBX2 and cPRC1. We performed CUT&RUN in FACS purified c-KIT(+) spermatogonia with antibodies against CBX2, RING1B, PHC2, BMI1 (PCGF4), H3K27me3, and H3K4me3. CBX2 was preferentially enriched at promoters of genes and its enrichment often spread >10kb (Fig. 4A and 4B). We obtained a conservative set of 708 promoters bound by CBX2 based upon enrichment of CBX2 using both CBX2 and HA antibodies (Fig. 4B). Notable examples of CBX2 targets identified in this set were *Foxc2*, *Gfra1*, and *Pax7* (Fig. 4A and Fig. S4A), key transcription factors and a cell surface receptor specifically expressed in spermatogonial stem cells and their immediate daughter cells (Aloisio et al., 2014; Hofmann et al., 2005; Wei *et al*., 2018). These genes are normally down-regulated as spermatogonia differentiate. On the contrary, genes robustly expressed in differentiating spermatogonia, such as *Sall4* and *c-Kit*, did not show enrichment of CBX2-cPRC1 (Fig. 4A and Fig. S4A). In general, transcripts associated with these 708 promoters had very low, median 0 FPKM expression in testis, based on a publicly available tissue-wide RNA-seq dataset. They had the highest expression from embryonic tissues, such as the embryonic brain or limbs (Fig. 4C) with enriched GO-terms often associated with PcG targets, such as ‘Homeobox’ (Fig. 4D). CUT&RUN analysis of RING1B, BMI1, and PHC2 all showed promoter-centered broad enrichment patterns that mirrored those of CBX2, showing that all four components of cPRC1 co-localized as expected (Fig. 4A and 4B). H3K27me3 also had expected overlapping enrichment patterns with CBX2 (Fig. 4A and 4B). The level of H3K4me3 enrichment anti-correlated with the levels of CBX2/cPRC1/H3K27me3 enrichment (Fig. 4B and Fig. S4B) on these genes, consistent with CBX2-cPRC1 bound genes having low transcriptional activity. We conclude that CBX2-PRC1 binds to genes that are normally down-regulated as spermatogonia differentiate and genes that were active in embryogenesis that need to be remain repressed.

**Figure 4.**
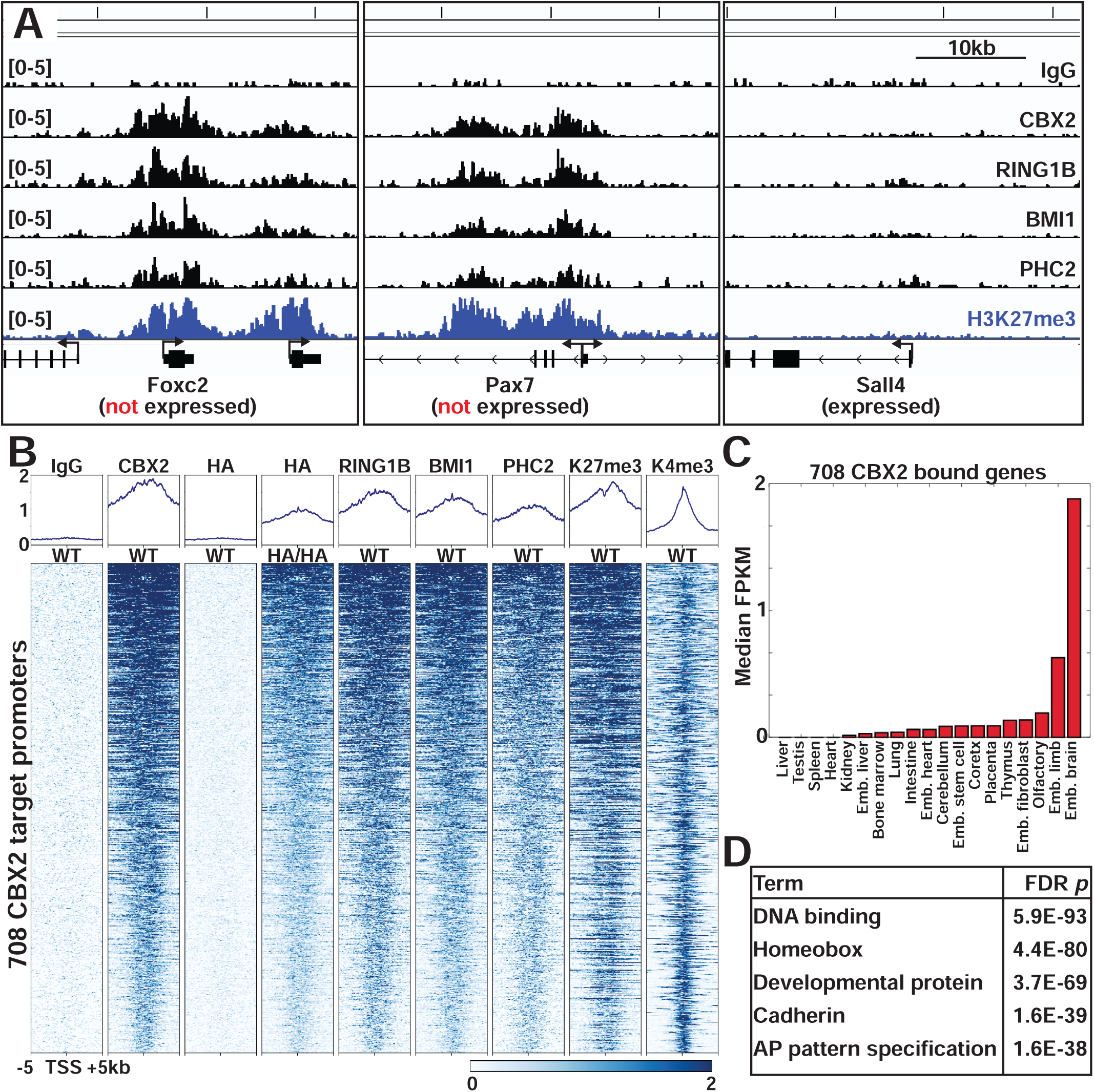
CBX2-cPRC1 binds genes expressed in spermatogonial stem cells and in embryonic tissues (A) Genome browser screenshots of CUT&RUN enrichment of IgG, CBX2, cPRC1 components (RING1B, BMI1, PHC2), and H3K27me3 from FACS sorted c-KIT(+) spermatogonia. Enrichment profiles at *Foxc2*, *Pax7* and *Sall4* genes are shown as representative examples. *Foxc2* and *Pax7* are highly expressed in spermatogonial stem cells and downregulated in c-KIT(+) differentiating spermatogonia.In contrast, *Sall4* is robustly expressed in c-KIT(+) spermatogonia. (B) Heatmaps showing CUT&RUN enrichment of CBX2, cPRC1 components, and associated histone modifications at 708 CBX2 target promoters. (C) Median expression levels of 708 CBX2 bound genes in 19 different mouse tissues at different developmental stages and in embryonic cell lines. Data are from Gene Expression Omnibus GSE29278 (Shen et al., 2012). (D) GO-term enrichment of 708 CBX2 bound genes.

### CBX2 is required for differentiation to CCND2(+) A1 spermatogonia

To test the function of CBX2 in spermatogenesis in adult animals, we used an inducible knock out (KO) strategy. We injected tamoxifen into a mouse line which expressed inducible CRE recombinase (*Rosa26-CRE^ERT2^*) and contained floxed *Cbx2* alleles (Fig. 5A and Fig. S5A). We chose an inducible KO strategy, as we aimed to assess immediate effects after inducing *Cbx2* deletion to minimize cellular compensation mechanisms. Even though *Rosa26* is a ubiquitous CRE driver, only spermatogonia express CBX2 in adult seminiferous tubules (Fig. 1). Both experimental (CRE+) and control (CRE-) animals received three times daily tamoxifen injections. Testes were collected at day 4, and changes in cell type distribution and gene expression were analyzed by single cell RNA sequencing after FACS purification of c-KIT(+), CBX2-expressing spermatogonia (Fig. 5A).

**Figure 5.**
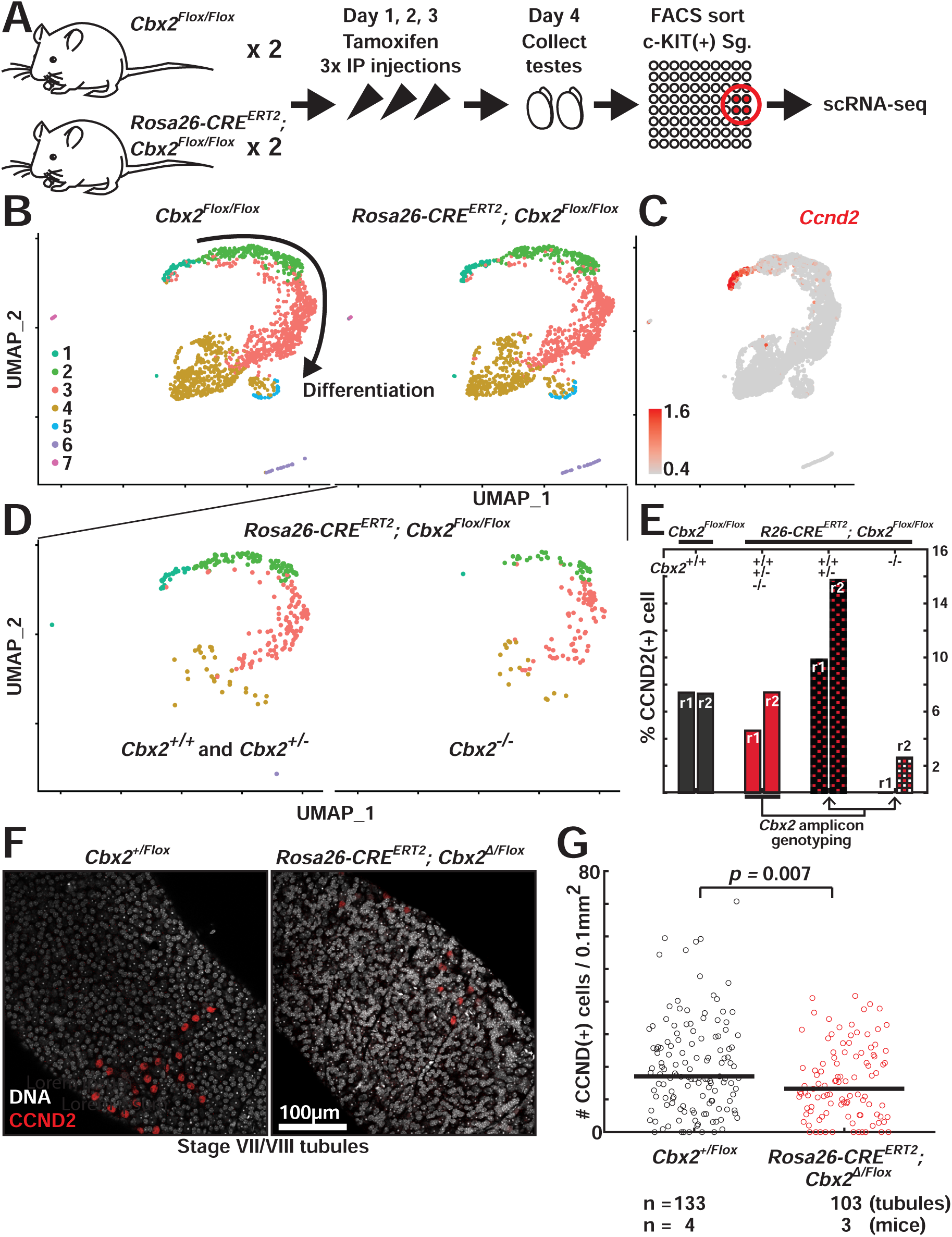
CBX2 is required for differentiation to CCND2(+) A1 spermatogonia (A) Schematic representation of experimental steps. (B) UMAP representation of a scRNA-seq comparison between FACS sorted c-KIT(+) spermatogonia from control (*Cbx2^Flox/Flox^*) and *Cbx2* inducible mutant (*Rosa26-CRE^ERT2^; Cbx2^Flox/Flox^*) animals. Each UMAP is a combined result of two replicates (two animals per condition). (C) Normalized expression level of *Ccnd2* is represented on the UMAP of all profiled cells. (D) Separate UMAP representation of *Cbx2^+/+^*, *^+/-^* and *Cbx2^-/-^* spermatogonia from *Cbx2* inducible mutant animals after *Cbx2* genotyping using *Cbx2*-specific amplicon sequencing. (E) Proportions of CCND2(+) cells (population #1 from Fig. 5B) among *Cbx2*-expressing populations (population #1, #2, #3) in different *Cbx2* genotype categories. Data from two replicates each (r1 and r2) for control and *Cbx2* inducible mutant animals are represented. (F) Immunofluorescence staining of whole testis tubules of control (*Cbx2^+/Flox^*) and *Cbx2* inducible mutant (*Rosa26-CRE^ERT2^; Cbx2^Δ/Flox^*) animals with CCND2 (red) antibody. Only stage VII/VIII tubules where CCND2(+) A1 spermatogonia reside are analyzed. Stage VII/VIII was identified by nuclear DNA staining patterns of characteristic pre-leptotene spermatocyte morphology and organization and by the presence of aligned condensed spermatid nuclei in the deeper plane. (G) Quantification of the number of CCND2(+) A1 spermatogonia per 0.1mm^2^ segments of stage VII/VIII tubule sections. Because A1 spermatogonia are not uniformly distributed on testis tubules, at least 30 different tubule segments were imaged per animal. *p* value was calculated by two-tailed t-test.

Single cell analyses of sorted c-KIT(+) cells identified four major populations of cells that encompassed >95% of all profiled cells (Fig. 5B, populations #1-4); the remaining <5% of cells were classified into three minor populations. As expected, all four populations expressed *c-Kit* (Fig. S5B). Population #1 was the most undifferentiated with specific expression of *Ccnd2*, which only marks A1 stage cells among differentiating spermatogonia (Fig. 5C and S2A) (Beumer et al., 2000). *Cbx2* was expressed in populations #1-3 but its expression was diminished in population #4, which started to express the second wave of *Stra8* as cells entered the pre-leptotene stage (Fig. S5B). Hence, we focused our subsequent analyses on populations #1-3 cells.

At first glance, scRNA-seq comparison of *Cbx2* mutant (CRE+) and control (CRE-) animals showed almost identical cellular distributions (Fig. 5B). This was surprising because ablation of PRC1, of which CBX2 is a key component in these cells, was previously shown to cause defects in the male germ cell line (Maezawa *et al*., 2017). However, The *Cbx2* mutant animals (CRE+) contained a mix of homozygous wild-type (*Cbx2^+/+^*), heterozygous (*Cbx2^+/-^*), and homozygous mutant (*Cbx2^-/-^*) cells, which we hypothesized obscured any differences between the bulk populations (Fig. 5B). We therefore separated cells obtained from the *Cbx2* mutant animals based on their genotypes, and the genotype-specific analysis revealed a distinct behavior of *Cbx2* homozygous mutant cells (Fig. 5D and 5E). We independently sequenced *Cbx2* amplicons generated from the aliquot of cell-barcoded cDNA derived from the samples prepared for single cell RNA-seq (Fig. S5A, S5C and S5D; see legends and methods). When the cell type distribution of *Cbx2* homozygous mutant cells (*Cbx2*^-/-^) were compared to *Cbx2* heterozygous mutant and homozygous wild-type cells within the same testes (*Cbx2^+/+^*, *Cbx2^+/-^* combined), we noticed a depletion of *Ccnd2*(+) A1 spermatogonia in *Cbx2^-/-^* cells (Fig. 5D and 5E). Among the CBX2- expressing population #1-3, while *Ccnd2*(+) A1 spermatogonia each occupied 7.9% and 7.8% of cells in two control animals (CRE-), they occupied only 0% and 2.8% *Cbx2^-/-^* cells isolated from the two *Cbx2* mutant (CRE+) animals (Fig. 5E). In contrast, A1 spermatogonia occupied 10.5% and 16.8% of *Cbx2^+/+, +/-^* cells isolated from the mutant. The increase in A1 spermatogonia *Cbx2^+/+, +/-^* cells in the mutant is consistent with the possibility that *Cbx2^+/+, +/-^* cells might compensate for the depletion of *Cbx2^-/-^* cells within the same testes (Fig. 5E). Only about 10∼20% of the total cells contained sufficient *Cbx2* reads to be confidently genotyped, which may have contributed to the variability of A1 spermatogonia proportion between replicates (r1 and r2) for genotyped cells. However, both replicates showed consistent trends of specific depletion of *Cbx2^-/-^* A1 spermatogonia. In addition, gene expression comparison between *Cbx2^+/+, +/-^* versus *Cbx2^-/-^* spermatogonia (population #1-3 combined) identified *Stra8* as the top downregulated gene in *Cbx2^-/-^* spermatogonia (Fig. S5E). *Stra8* (*<u>St</u>imulated by <u>r</u>etinoic <u>a</u>cid gene 8*) is a gene upregulated in A1 spermatogonia by retinoic acid stimulation (Fig. 2A and S2A). Lower *Stra8* level in this population is consistent with the depletion of A1 spermatogonia in *Cbx2^-/-^*cells. We conclude based upon this single cell *Cbx2* genotyping of mosaic mutant testis that CBX2 plays a key role in the production of A1 spermatogonia.

We confirmed the depletion of CCND2(+) A1 spermatogonia by independent immunostaining experiments on wholemount tubules. We could not confidently tell *Cbx2* genotypes by immunostaining due to limited antibody sensitivity, so we performed experiments by inducing as complete a homozygous null *Cbx2* mutation as possible using a mouse line where one allele of *Cbx2* was already deleted (see methods). Quantification of CCND2(+) cells at stage VII/VIII tubules by wholemount immunostaining showed a significant decrease of CCND2(+) cells in *Cbx2* mutant compared to the control animals (Fig. 5F and 5G). The number of pre-leptotene spermatocytes in the same stage VII/VIII tubules did not show significant differences (Fig. S5F), suggesting CBX2 function is specific to the production of CCND2(+) A1 spermatogonia. Notably, the phenotype of CBX2 depletion on A1 spermatogonia coincides with the strong upregulation of CBX2 expression at the A1 stage in wild-type animals (Fig. 2C and 2D). We did not detect a significant increase in the number of apoptotic cells in the stage VI to VIII tubules where Aaln-to-A1 transition happens (Fig. S5G and S5H), suggesting that CBX2 function is not likely to be required for the survival of germ cells. We conclude that CBX2 is required for appropriate differentiation, specifically for the transition to the differentiating spermatogonia.

### The CBX2 CaPS domain is required for germ cell maintenance

CBX2 can both compact nucleosomal arrays and phase separate, activities that both rely on the positive charge of the central ‘CaPS’ (Compaction and Phase Separation) domain (Fig. S1A). We examined the hypothesis that this domain might be important for CBX2 function in spermatogenesis by using a knock-in mouse line in which the endogenous CaPS domain of both CBX2 alleles was mutated. The replacement of 23 positively charged amino acids (Lys, Arg) of CBX2 with a neutral amino acid (Ala, hereafter *Cbx2^23KRA^* mutant, Fig. S1A) in the CaPS domain diminished polynucleosome compaction (Grau *et al*., 2011) and phase separation activities (Plys *et al*., 2019) *in vitro*, and resulted in posterior transformation of axial skeletons *in vivo* (Lau *et al*., 2017). Using the CaPS domain mutant, we tested whether this functional domain of CBX2 played a role in adult tissue homeostasis during spermatogenesis. The CBX2^23KRA^ mutant does not disrupt other domains needed for chromatin targeting and for cPRC1 formation.

*Cbx2^23KRA/23KRA^* mice showed progressive defects in male germ cell maintenance as mice aged. Both heterozygous and homozygous *Cbx2^23KRA/23KRA^*mutant mice showed largely normal development and fertility albeit having axial skeletal defects (Lau, 2016). Consistent with the results from mESCs (Jaensch *et al*., 2021) and MEFs (Isono *et al*., 2013) that factors in addition to CaPS also are able to drive Polycomb body formation, spermatogonia from wild-type and *Cbx2^23KRA/23KRA^* mice did not show visible differences in CBX2-Polycomb puncta (Fig. S6A). While *Cbx2^23KRA/+^*and *Cbx2^23KRA/23KRA^* mice had similar body weights when compared to wild-type siblings up to ∼40 weeks of age (Fig. 6A), the *Cbx2^23KRA/23KRA^*mutant mice had lower testis weights than their wild-type siblings as the mice got older, especially over 30 weeks (Fig. 6B and 6C). This result indicates a progressive defect in spermatogenesis as mice aged. Histological examination of testis sections of old (>30 weeks) *Cbx2^23KRA/23KRA^* mutant mice revealed clusters of defective tubules that were devoid of all germ cells and only had Sertoli cells at the basement membrane (Fig. 6D, 6E, and 6G). Five percent of tubules (median value) were devoid of germ cells in homozygous *Cbx2^23KRA/23KRA^* mutant mice, which was significantly higher than in wild-type siblings, which rarely showed germ cell-less tubules (Fig. 6F). Some tubules exhibited a ‘missing generation’ phenotype (Lovasco et al., 2015). These tubules lack spermatocytes (Fig. 6H and 6I), indicating that SSCs did not differentiate in one of previous differentiation cycles, potentially to preserve their declining numbers or due to problems in differentiation.

**Figure 6.**
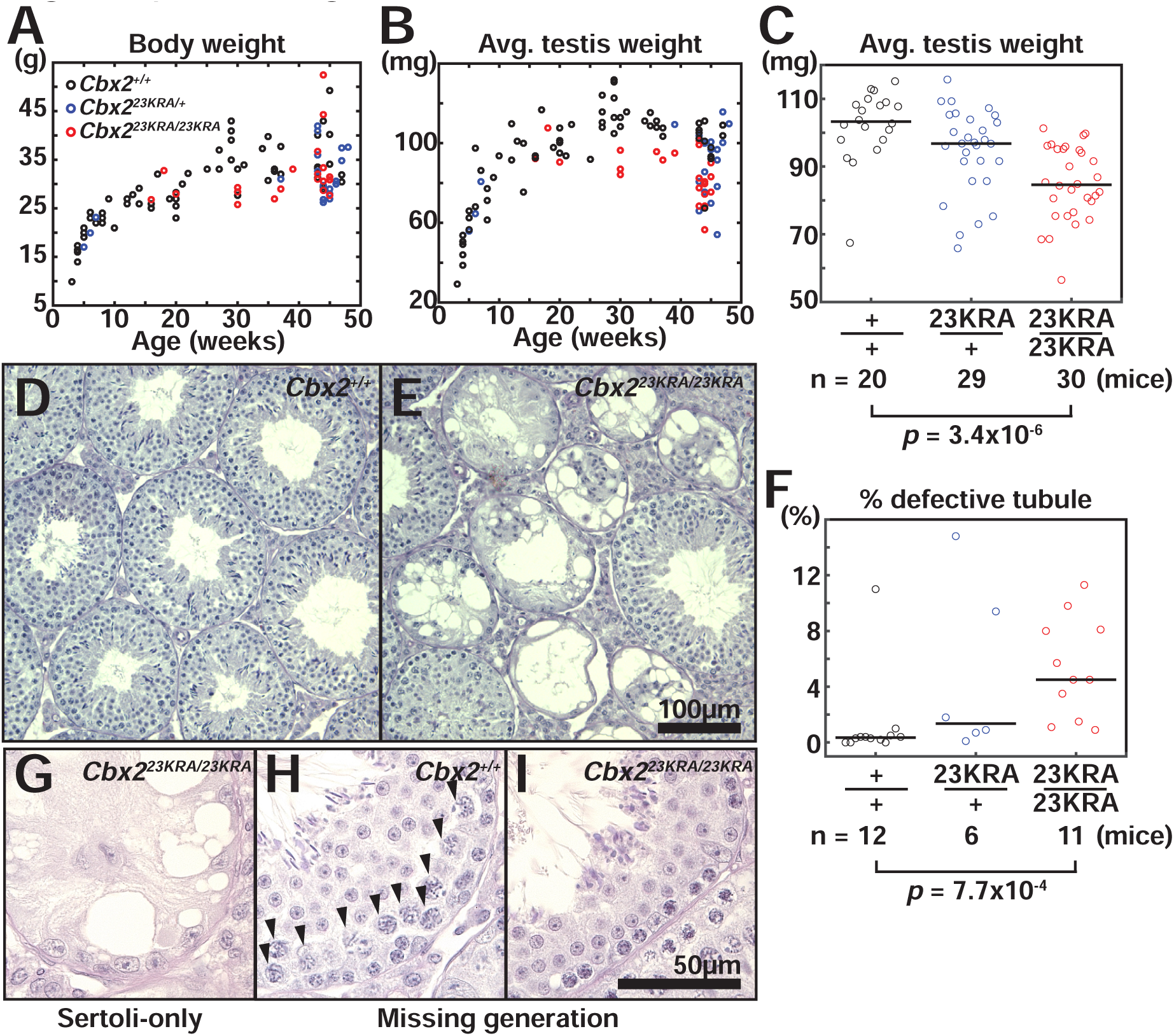
Spermatogenesis defects in CBX2 CaPS domain mutant mice (A and B) (A) Body and (B) average testis weight of wild-type, *Cbx2^23KRA/+^*, and *Cbx2^23KRA/23KRA^* mutant mice at the time of euthanasia. (C) Box plot of average testis weights of aged (more than 30 weeks old) mice. The *p* value is based on Mann-Whitney U test. (D and E) Representative images of Hematoxylin-PAS-stained testis sections of sibling (D) wild-type and (E) *Cbx2^23KRA/23KRA^* mutant mice. (F) Quantification of percentage of tubules with defects in germ cell maintenance in wild-type, *Cbx2^23KRA/+^* and *Cbx2^23KRA/23KRA^* mutant mice. The *p* value is based on Mann-Whitney U test. (G-I) Representative images of Hematoxylin-PAS-stained testis sections showing (G) Sertoli cell-only, (H and I) missing generation phenotypes, (I) lacking spermatocytes (arrowheads in H).

It was notable that only a fraction of tubules showed complete loss of germ cells, while adjacent tubules seemed to maintain normal spermatogenesis (Fig. 6E). To probe if there are subtle defects in the seemingly normal tubules in the *Cbx2^23KRA/23KRA^* mice, we counted the numbers of A spermatogonia, pre-leptotene and zygotene spermatocytes in stage VII/VIII and XII tubules using histological sections from old (>30 weeks) mice (Fig. S6B and S6C). In all cases, there were no statistically significant differences between wild-type and *Cbx2^23KRA/23KRA^* mutants in the number of germ cells (Fig. S6D-G). However, the number of A_undiff_ and A1 spermatogonia per tubule section showed a slight decrease in the median number, with *Cbx2^23KRA/23KRA^* mice having more tubule sections without any spermatogonia (Fig. S6D). No such decrease was observed in other cell types in the lineage (Fig. S6E-G). These data indicate that the regulation of the ratio of spermatogonia to spermatocytes is mostly normal except for a minor imbalance in A_undiff_/A1 spermatogonia distribution. This imbalance might, over time, contribute to defects in spermatogenesis. In addition, this result suggests that disruption of the CaPS domain in CBX2 likely plays a similar role as to the deletion of CBX2, which showed a significant depletion of A1 spermatogonia. Notably, there were no significant differences in binding of CBX2 or enrichment of H3K27me3 modifications at 708 CBX2 bound genes between wild-type and *Cbx2^23KRA/23KRA^* c-KIT(+) spermatogonia by CUT&RUN (Fig. S6H and S6I). Thus, for long term tissue homeostasis, a non-enzymatic activity of CBX2-cPRC1 is critical for preventing stochastic loss of germ cells, and hence, maintenance of spermatogonial stem cells.

## Discussion

The core function of the PcG family of complexes is to maintain gene expression patterns across the numerous cell divisions required to form and sustain an animal. Our characterization of the role for the CBX2 subunit and for its main functional domain during spermatogenesis advances the understanding of how that cell fate regulation is accomplished in a specific lineage in an animal. CBX2 is upregulated in a cell type-specific manner as spermatogonial stem cells initiate differentiation, and it forms nuclear condensates with its target genes. Single cell comparison of *Cbx2* wild-type and mutant cells from the same mosaic mutant testes revealed that CBX2 function is critical at an early stage of differentiation corresponding to when CBX2 expression was upregulated. The domain responsible for compaction and phase separation of CBX2 was required specifically for male germ cell maintenance (Fig. 7). We propose that the ability of CBX2-cPRC1 to alter chromatin structure is critical for a specific step in spermatogenesis (the formation of CCND2(+) A1 spermatogonia) and for maintenance of a key adult stem cell lineage.

**Figure 7.**
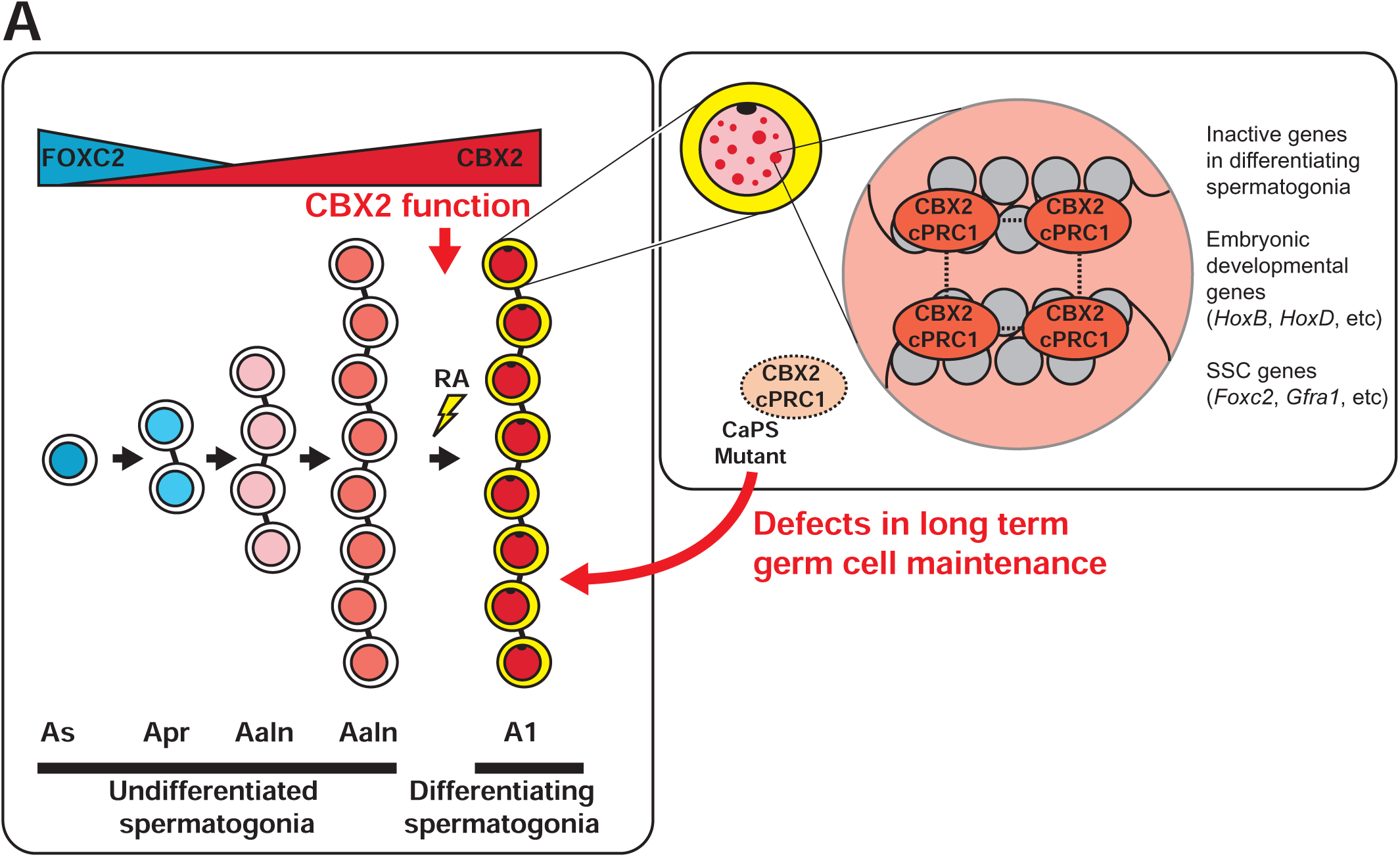
A model of CBX2 function at the exit from the SSC state CBX2 expression is upregulated as A single or A paired spermatogonia differentiate to A aligned spermatogonia. CBX2 expression further increases at A1 spermatogonia when the cells irreversibly commit to differentiation. CBX2 function is required specifically for the production of A1 spermatogonia. In differentiating spermatogonia, CBX2 forms Polycomb bodies that encompass its target genes that need to remain repressed. Disruption of CaPS domain of CBX2 results in defects in the long-term maintenance of germ cells.

Some PcG proteins are specifically expressed and required in certain adult stem cell types (Flora *et al*., 2021). In contrast, we found that CBX2 expression was relatively low in spermatogonial stem cells, was upregulated as stem cells initiate differentiation, and was necessary for proper amounts of A1 spermatogonia. Therefore, in the male germ cell lineage, CBX2-cPRC1 is not an exclusive regulator of stem cells but instead is more active in committed progenitor cells. This CBX2 expression pattern is reminiscent of its expression during differentiation in cultured mouse embryonic stem cells (mESCs), in which CBX2 is upregulated as mESCs differentiate into embryoid bodies or neural progenitor cells (Morey et al., 2012; O’Loghlen et al., 2012). In addition, *Cbx2* was shown to be upregulated in differentiating human bone marrow cells, but not in immature cells (Lessard et al., 1998). Thus, CBX2 expression is associated with lineage commitment in diverse developmental contexts. Here, we show that CBX2 is necessary for the proper differentiation of cells during spermatogenesis specifically at the stage when it is upregulated, not at the stem cell stage.

Why is the number of A1 spermatogonia reduced in *Cbx2^-/-^*lineages? There was no significant increase of apoptosis in *Cbx2* mutant germ cells (Fig. S5G and S5H), indicating that CBX2 function is needed either for A aligned-to-A1 differentiation or production of sufficient precursor A aligned spermatogonia. A known function of PcG complexes is the repression of cell cycle inhibitor genes (Jacobs et al., 1999). It is possible that A aligned spermatogonia in the *Cbx2* mutant testes might divide less due to incomplete repression of the cell cycle inhibitors. Alternatively, it is possible SSCs do not stably exit from their stem cell program to become A aligned spermatogonia. Undifferentiated spermatogonia normally decrease the expression of SSC-specific genes, such as FOXC2, as they become differentiation-competent (RARg+, retinoic acid-responsive) A aligned spermatogonia (Ikami *et al*., 2015; Wei *et al*., 2018). Because CBX2 directly binds SSC-specific genes in differentiating spermatogonia, it might be that *Cbx2* mutant A aligned spermatogonia cannot efficiently maintain the repression of SSC genes. While the exact mechanism remains to be elucidated, the key conclusion is that CBX2 is not a general factor required for all dividing cells, but rather it regulates a specific developmental transition step.

Phase separation has been proposed to play a key role in maintenance of repressive states based upon studies of the heterochromatic protein HP1 (Larson et al., 2017; Strom et al., 2017) and based upon previous analysis of CBX2 and PcG function (Plys *et al*., 2019; Tatavosian *et al*., 2019). There is scant information concerning the extent to which phase separation occurs in living adult animals and the extent to which it might contribute to maintenance of stably differentiated tissues. We show that in testes there is overlap of puncta containing CBX2-PRC1 and *HoxD* and *HoxB* target genes. Therefore, these puncta, previously shown to be present in embryos (Grimaud *et al*., 2006) and cultured cells (Isono *et al*., 2013; Satijn *et al*., 1997; Saurin *et al*., 1998), also exist in an adult animal tissue and are co-resident with regulated genes. Although the *HoxD* locus did not overlap with CBX2 or RING1B signal in ∼30% of the images, this degree of non-overlap is similar to what was observed in MEFs, showing ∼20% of non-overlap between PCGF2 (MEL-18) and *HoxB* loci (Isono *et al*., 2013). This degree of non-overlap might be technical as heat applied to denature the DNA duplex disrupts immunostaining signal patterns and intensity. Alternatively, it might be that in certain cell cycle phases, for example S-phase, Polycomb target loci may temporarily be displaced from Polycomb bodies.

Aged mice with a mutation in the domain required *in vitro* for both phase separation and chromatin compaction (‘CaPS’ domain) developed sporadic empty tubules in the testes. These mutant mice still had visible puncta of CBX2-PRC1 (Fig. S6A). This was anticipated based upon previous data in cell culture (Jaensch *et al*., 2021) and the known phase separation and oligomerization characteristics of other cPRC1 components, especially the PHC proteins (Isono *et al*., 2013; Seif *et al*., 2020). The incomplete penetrance of the empty tubule phenotype mirrors previous studies of mutant PRC1 components, which also show incomplete penetrance (Kim and Kingston, 2022; Takada et al., 2007). This might be due to redundant activities of these components and their paralogs. For example, *Cbx4* and *Cbx8* were expressed in differentiating spermatogonia (Fig. S2A) and were upregulated in *Cbx2* KO embryos (Fig. S1C). Even though Polycomb bodies are still present in the *Cbx2* CaPS mutant testes, the biophysical properties of these condensates might differ from those formed in wild-type mice in a manner that decreases the effectiveness of cPRC1 in driving memory of cell fate but does not eliminate memory.

In aged *Cbx2* CaPS domain mutant mice, some tubules show complete loss of germ cells, implying loss of SSCs. As CBX2 was required for commitment to A1 cells when acutely depleted, it is counterintuitive that the precursor SSCs are also lost in the *Cbx2* CaPS mutant testes. However, the depletion of germ cells in *Cbx2* CaPS mutant testes only becomes noticeable after 30 weeks (Fig. 6), which corresponds to up to 20 cycles of new A1 spermatogonia production events. Disruption of the initial phase of differentiation (the production of A1 spermatogonia) might generate a feedback loop in which, over time, accumulation of stress could cause some tubules to lose SSCs. Similarly, the knock out of another PcG gene *Eed* did not alter the initial hematopoietic stem cell (HSC) establishment in fetal liver but showed defects in their differentiation and ultimately led to their depletion due to exhaustion (Xie *et al*., 2014).

Another noticeable feature of the *Cbx2* CaPS mutant is that some tubules show complete loss of germ cells, while neighboring ones maintain seemingly normal spermatogenesis. Total output of sperm is regulated in part by over-production of undifferentiated spermatogonia followed by regulated proliferation, migration and apoptosis of differentiating spermatogonia (de Rooij and Russell, 2000; Yoshida et al., 2007). Therefore, even if there is a reduction in the number of SSCs, the remaining SSCs in principle can support normal spermatogenesis until the number falls below critical levels within a specific tubule. This all-or-nothing phenotype was also observed in mutants affecting SSC maintenance, such as *Zbtb16* (Buaas et al., 2004; Costoya et al., 2004). The stark difference in the penetrance between tubules might arise due to the combined impacts of the robust nature of spermatogenesis and the stochastic nature of PcG gene mutants.

## Conclusion

Like other epigenetic regulators whose normal function is to ensure proper development and differentiation, PcG genes are frequently dysregulated in human cancers and aging. To utilize chromatin pathways to reverse such conditions, chemical inhibitors of PcG function have been developed. However, to provide effective intervention, it is essential to understand how PcG dysfunction is involved in complex processes, such as the decline of tissue function. We demonstrate that CBX2 acts at the exit from the spermatogonial stem cell state for the commitment of differentiation, and the domain required for compaction and phase separation is critical for the long-term maintenance of the lineage. Thus, the structural activity of cPRC1 driven by CBX2 is crucial for proper developmental transitions in the male germ line and for maintenance of the complete germ line machinery during aging. It will be important to investigate if cPRC1 plays similar roles in cell fate regulation in other prolific adult stem cell lineages.

## Limitation of the study

An important remaining question concerns whether there are changes in chromatin structure that occur *in vivo* following disrupting CBX2 and, if so, what the nature of those changes might be. There are several challenges in addressing this problem. First, the chromatin state of Polycomb target regions is difficult to measure with accessibility assays, including ATAC-seq, because sequencing coverage is highly biased to ’open’ chromatin, such as active enhancers and promoters. Second, it is challenging to isolate spermatogonia undergoing the Aaligned-to-A1 transition, where CBX2 function is critical, using cell surface markers. Third, potential compensation by paralogs, such as CBX4 or CBX8 can also dampen the effect. In future studies, cell culture models can be applied for molecular analyses. Spermatogonial stem cells can be cultured *in vitro* (Kanatsu-Shinohara et al., 2003), and the first step of differentiation can be modeled by retinoic acid stimulation. Using more acute and complete deletion of *Cbx2* from SSCs derived from the floxed *Cbx2* mice, chromatin structure can be assessed with established methods, such as MACC (Mieczkowski et al., 2016), from more homogenous and abundant sources of cells.

## Materials and methods

### Mice

All animal procedures were conducted in accordance with protocols approved by the Massachusetts General Hospital Institutional Animal Care and Use Committee (IACUC, Protocol# 2015N000072), and animals cared for according to the requirements of the National Research Council’s Guide for the Care and Use of Laboratory Animals.

The floxed *Cbx2^Flox^* mouse line was generated from the parental ’targeted trap’ line (Cbx2tm1a(KOMP)Wtsi, in short *Cbx2^tm1a^*), that was originally produced by the Knockout Mouse Project (KOMP) (Skarnes et al., 2011). Aliquots of cryopreserved sperm (*Cbx2^tm1a^*) were obtained from the UC Davis Mouse Biology Program KOMP, and *in vitro* fertilization was performed by the Charles River Laboratories. Rederived *Cbx2^tm1a^*females were crossed to male mice expressing FLP recombinase (C57BL/6N-Tg(CAG-Flpo)1Afst/Mmucd, Stock# 036512-UCD) to excise the LacZ reporter trap flanked by FRT sites to make ’conditional-ready’, *Cbx2^Flox^*(or denoted as *Cbx2^tm1c^* per European Mouse Mutant Cell Repository nomenclature) mice. To produce an inducible *Cbx2* deletion line, *Cbx2^Flox^* mice were crossed to the mice expressing CRE recombinase responsive to Tamoxifen (R26-CreERT2, full genotype: B6.129-Gt(ROSA)26Sortm1(cre/ERT2)Tyj/J, Jackson Laboratory stock number: 008463). Inducible *Cbx2* deletion line with one deleted *Cbx2* allele (*R26-CreERT2*; *Cbx2^Δ/Flox^*) was produced by crossing *Cbx2^Δ/+^* males to *R26-CreERT2/+*; *Cbx2^Flox/Flox^* females. *Cbx2^Flox^* and its derivative lines were backcrossed to C57BL/6J mice more than 5 times. *Cbx2^23KRA^* strain (Lau *et al*., 2017) is available through the Mutant Mouse Resource and Research Center (MMRRC, B6J.Cg-Cbx2tm1.1Rek/Mmnc, Stock# 050534-UNC). *Cbx2^Δ^* strain has a deletion (mm10, chr11:119,028,119-119,028,164) by CRISPR/Cas9 that introduces a premature stop codon after amino acid 171. For the *Cbx2^23KRA^*strain, only non-stud males were used for aged phenotyping experiments to minimize the potential influence of the presence of females on spermatogenesis (Schmidt et al., 2009). *Cbx2^23KRA^* and *Cbx2^Δ^* strains were backcrossed to C57BL/6J more than 8 times. All C57BL/6J mice used for backcross were obtained from the Jackson Laboratory (Stock# 000664). PCR primers for genotyping are listed in ‘Supplemental table 1’.

### Tamoxifen administration

Tamoxifen (Sigma-Aldrich, Cat# T5648) was dissolved in corn oil (Spectrum Chemical, Cat# CO136) to a final 20 mg/mL by incubating the mixture at 37 °C overnight while rotating. Dissolved tamoxifen was filtered and aliquots for daily use were stored in sterile tubes and kept at 4 °C for up to 4 days. For the *Cbx2* amplicon single cell RNA-seq experiments, 150 µL of warmed-up (room temperature) tamoxifen-corn oil was injected intraperitoneally daily for 3 consecutive days. For CCND2 and cleaved-Caspase 3 immunostaining experiments, 200 µL of warmed-up tamoxifen-corn oil was injected intraperitoneally daily for 4 consecutive days to make as complete *Cbx2* mutant as possible.

### Generation of Cbx2^HA^ strain

To knock-in HA epitope tags at the N-terminus of *Cbx2* gene, guide RNA (Synthego), single stranded repair oligo (Genewiz), and Cas9 enzyme (Alt-R S.p. Cas9 Nuclease V3, Integrated DNA Technologies, Cat# 1081058) were electroporated into B6 mouse zygotes by the Genome Modification Facility, Harvard University. DNA sequences are provided in ‘Supplemental table 1’. Predicted 295 bp (232 bp for WT) knocked-in DNA fragment was PCR-amplified from genomic DNA of targeted pups. PCR fragments from 4 pups (out of 13) with the expected knock-in size were deep-sequenced by MGH CCIB DNA core (CRISPR Amplicon sequencing). The following 63bp sequence was knocked in right after the ATG start codon (TATCCATACGATGTTCCTGACTATGCGGGCTATCCCTATGACGTCCCGGACTATGCA GGATCC, inserted right after the position chr11:119023111, mm10). One *<u>Gly linker</u>* was placed between HA sequences, and one *<u>GlySer linker</u>* was placed between 2xHA and CBX2 protein (YPYDVPDYA*<u>G</u>*YPYDVPDYA*<u>GS</u>*). Three (2 HA/deletion males and 1 HA/HA female) mice contained the correctly targeted allele. Two founder males sired progenies. Homozygous *Cbx2^HA/HA^* males were produced by crosses between the progeny of the two different founders to minimize potential off-target mutations becoming homozygous. Targeted strains were backcrossed to C57BL/6J mice for 3 times at the time of immunostaining and CUT&RUN experiments.

### Testis dissociation to obtain single cells

Testes were obtained and tunica albuginea membrane was removed. Released seminiferous tubules were gently separated in PBS. Roughly separated tubules were incubated in collagenase type IV in DMEM (5 mL for 1 testis pair, 1 mg/mL, STEMCELL Technologies Inc., Cat# 07909) at 37 °C while rotating. After an initial 10-minute incubation, the collagenase solution with released interstitial cells was removed, and the same volume of fresh collagenase solution was added to the seminiferous tubules. Tubules were further dissociated by incubating at 37 °C for another 10 minutes with occasional stirring and pipetting up and down. Dissociated tubules were centrifuged at 300 × g for 3 minutes. The supernatant was removed and trypsin (Thermo Fisher Scientific, Cat# 12605010) solution was added to make a single cell suspension. After 5 minutes of Trypsin incubation at 37 °C with occasional mixing, dissociated cells were passed through 40 µm-wide cell strainers. One volume of 4 °C DMEM (Thermo Fisher Scientific, Cat# 11960044) with 10% fetal bovine serum (Sigma-Aldrich, F2442) was added and the single cell suspension was centrifuged at 400 × g for 6 minutes. Cell pellets were resuspended in 4 °C DMEM with 10% FBS.

### Fluorescence activated cell sorting (FACS)

Cells were resuspended in PBS at 4 °C at ∼1×10^7^/mL concentration. Cells were first stained with a cell viability dye Zombie Green (BioLegend, Cat# 423111) for 20 minutes at room temperature. After one wash with PBS (+0.5% (w/v) BSA), cells were incubated with cell surface antibodies in PBS (+0.1% (w/v) BSA) at 4 °C for 30 minutes. Antibody product and dilution information are listed in the ‘Key resources table’. After antibody incubation, cells were washed with 50 mL 4 °C PBS once, and then resuspended in PBS (+0.1% (w/v) BSA) at ∼1×10^7^/mL concentration. FACS was performed with BD FACS Melody.

### Preparation of cell and tissue lysates for immunoblotting

Dissociated single cells or tissue material were dissolved in SDS tissue lysis buffer (20 mM Tris-HCl, pH 8.0, 135 mM NaCl, 10% glycerol, 1% Igepal, 1% SDS, 5 mM EDTA). 1 µL Benzonase (25KU, Millipore, Cat# 71205) was added per 200 µL of lysis buffer. Tissue pieces were homogenized by passing through 23G and 25G needles consecutively at least 10 times. Cell and tissue lysates were incubated for about 15 minutes at room temperature until the lysate was no longer viscous. If the lysates were still viscous, they were sonicated with Bioruptor at ‘High’ setting for 4 minutes (30 sec on/30 sec off). Whole cell or tissue lysates were used for immunoblotting without further separation by centrifugation.

### Western blot

Cell or tissue lysates (10 to 20 µg) were mixed with 5X western loading buffer and incubated at 95 °C for 5 minutes. Samples were then loaded to 4-20% polyacrylamide gels in western running buffer (25 mM Tris, 192 mM Glycine, 0.1% SDS (w/v)) at constant ∼120 V for 2∼3 hours. After separation, proteins were transferred to PVDF membrane (Fisher Scientific, Cat# IPFL00010) in western transfer buffer (25 mM Tris, 192 mM Glycine, 20% methanol(v/v)) at a constant 45 mA overnight for a total 400 volt-hours. After confirming the transfer with Ponceau staining, membranes were washed once with wash buffer (PBS with 0.1% Tween 20) and incubated in a blocking buffer (PBS with 0.1% Tween 20, 5% (w/v) skimmed milk) for 30 minutes at room temperature. Membranes were incubated with primary antibodies with desired concentrations in the incubation buffer (PBS with 0.1% Tween 20, 2% (w/v) skimmed milk) overnight at 4 °C. The concentrations of antibodies and their sources are listed in the ‘Key resources table’. Membranes were washed three times in the wash buffer (PBS with 0.1% Tween 20) and incubated with secondary antibodies conjugated with HRP in the incubation buffer overnight at 4 °C. Membranes were washed three times and developed with ECL solution (Thermo Scientific, Cat# 1859698, 1859701).

### Tissue preparations for RNA in situ hybridization, immunostaining and histology

Dissected testes and epididymides were fixed in >10 mL of 4% formaldehyde in PBS (for immunostaining or *in situ* hybridization) at 4 °C or in Bouin’s fixative (for Hematoxylin/PAS staining) at room temperature overnight with rocking. Testes were cut in half the next day and fixed with the same fixatives for two more hours. After washing with water, testes and epididymides were dehydrated with successive incubation in 30%, 50% and 70% ethanol for 1 hour each. Fixed and dehydrated testes and epididymides were paraffin-embedded and cut in 5 µm thickness for subsequent experiments.

### Immunostaining of testis sections

Cut sections on microscope slides were baked at 60 °C for one hour in an oven. The paraffin was removed by incubating the sections in Xylene for 10 minutes 3 times, and residual Xylene was washed away by incubating in 100% ethanol for 5 minutes 3 times. Samples were subsequently rehydrated by incubating in 95%, 85%, and 70% ethanol for 3 minutes each and rinsed with >4 L of water. Deparaffinated samples were antigen retrieved by incubating in boiling retrieval buffer (10mM Sodium Citrate at pH 6.0, 0.05% (v/v) Tween-20) for 10 minutes. Boiled slides were cooled and washed with >4L water. Samples were incubated with a blocking buffer (3% BSA (w/v), 0.05% Triton X-100 (v/v), PBS) for 30 minutes at room temperature. Samples were incubated with primary antibodies in a blocking buffer overnight at 4 °C, washed three times with PBST (0.05% Triton X-100 (v/v) PBS). Primary antibody information is listed in the ‘Key resources table’. Samples were then incubated with secondary antibodies in a blocking buffer overnight at 4 °C, washed four times with PBST. The second wash was done with PBST with HOECHST 33342 (final 1 µg/mL, Life Technologies, Cat# H3570). Both primary and secondary antibody incubations were done in a humidified chamber. Coverslips were put on top of sections with mounting media (Vector Laboratories, Vectashield, H-1000) and sealed with nail polish (Fisher Scientific, Cat# 72180). Slides were imaged with the Nikon 90i Eclipse epifluorescence microscope equipped with an ORCA-ER-1394 CCD camera (Hamamatsu).

### Immunostaining of testis tubules

Testis tubules were gently separated in 4 °C PBS. Dissociated tubules were fixed in 4% formaldehyde in PBS for 15 minutes with occasional rocking at room temperature. Fixed tubules were then quickly washed with PBS twice, incubated in 100 mM Tris pH 8.0 for 10 minutes to quench unreacted formaldehyde, and washed twice with PBS. Fixed tubules were transferred to 10 cm Petri dishes in PBST (0.1% Triton X-100). Tubules were cut into ∼1 mm length pieces under a dissection microscope. When imaging stage VI-VIII tubules, ’dark zone’ tubules were collected because in stage VII, spermatids move towards the lumen showing characteristic dark patterns in the middle of tubules (Kotaja et al., 2004). More than 30 pieces of ∼1 mm length tubules were transferred to test tubes using wide bore tips and blocked in PBST BSA (0.05% Triton X-100, 3% BSA(w/v)) for >15 minutes. Tubules were then incubated with primary antibodies for two nights at 4 °C, washed three times with PBST (0.05% Triton X-100), then incubated with fluorophore-conjugated secondary antibodies overnight, then washed four times with PBST (0.05% Triton X-100). For the second wash, HOECHST 33342 (final 1 µg/mL, Life Technologies, Cat# H3570) was added to label nuclei. Washed tubules were mounted on glass slides, arranged on the slide in a non-overlapping manner, and mounted with mounting media (Vector Laboratories, Vectashield, H-1000). Tubules were imaged with Leica TCS SP5 confocal microscope with at least two times line averaging for 1024×1024 pixels. All settings, including laser intensity, gain and offset, and pinhole size, kept constant for the samples to be compared. Pixel intensity was measured by manually selecting spermatogonial nuclei with ’Oval selections’ (6 µm diameter) in Fiji (Schindelin et al., 2012).

### Immunostaining of dissociated cells on coverslips

Coverslips (12mm circles, thickness 1, Fisher Scientific, Cat# 72231-01) were incubated with Poly-D-Lysine (R&D Systems, Cat# 3439-100-01) for >5 minutes prior to the cell attachment. 1mL of dissociated cells from testes (>1×10^5^ cells per well) in PBS were applied on the coverslips in a 24-well plate. The plate was briefly centrifuged for 1 minute at 80 × g. The attached cells were fixed with 1% formaldehyde in PBS (v/v) for 15 minutes at room temperature. After fixation, coverslips were washed with PBS twice and PBST once. Coverslips were blocked and incubated with antibodies in 24-well plates as specified in immunostaining of testis sections. Mounted coverslips were imaged with Leica TCS SP5 confocal microscope with at least two times line averaging for 512×512 pixels.

### CBX2 antibodies

N-terminal FLAG epitope-tagged near full-length CBX2 (FLAG-CBX2-ΔCbox, aa1-485) protein was used as an antigen. C-terminal Cbox was deleted to prevent CBX2 from degradation when overexpressed without its binding partner, RING1B. CBX2 protein was expressed in Sf9 cells using the baculovirus system and affinity purified from nuclear extracts using agarose beads coupled with the M2 anti-FLAG antibody (Sigma-Alrich, Cat# A2220). Purified CBX2 protein (1 mg, 1 mg/mL) was each injected into two rabbits for initial immunization. Subsequently, 0.2 mg (0.5 mg/mL) of purified CBX2 protein was injected for boosts at day 14, 21, 49, and 77. Final antisera were obtained on day 91. Antibodies against CBX2 were affinity purified using the Affi-gel (Bio-Rad, Affi-Gel 10, Cat# 1536099)-coupled FLAG-CBX2-ΔCBox, concentrated and stored in PBS with 10% glycerol. All rabbit experiments were done in Cocalico Biologicals with USDA research license and animal welfare assurance with NIH (http://www.cocalicobiologicals.com/antibodies.html).

### RNA in situ hybridization

*Cbx2 in situ* hybridization on testis sections was performed using the RNAScope (Advanced Cell Diagnostics, Cat# 322300) based on manufacturer’s protocol. Antigen retrieval was performed for 10 minutes in a boiling antigen retrieval buffer. *Cbx2* probes are targeting 541-1606 region of transcript NM_007623.3. Images were acquired with Leica DM5000B microscope.

### Generation of DNA-FISH probes

*HoxD* (mm10, chr2:74653050-74762090) and gene desert (mm10, chr11:37675694-38150801) probes were Alexa 647 labeled oligopaint probes used in the previous study (Kundu *et al*., 2017), and generated based on the published protocol (Beliveau et al., 2012). *HoxB* (mm10, chr11:96349820-96368979) was visualized using SABER-FISH, and the probes were generated based on the published protocol (Kishi et al., 2019) with the following modifications. The primer exchange reaction (PER) was performed for at least 3 hours. The resulting probe concatemer length was verified using agarose gel electrophoresis with EtBr after making the concatemer part of the probes double stranded by hybridizing non-fluorescent imager oligos. SABER-FISH oligo sequences are listed in ‘Supplementary table 1’.

### Co-immunofluorescence staining DNA-FISH

FACS-sorted germ cells were attached to the coverslips and immunostained with desired antibodies with the staining protocol described in the ‘*Immunostaining of dissociated cells on coverslips*’ section. After staining, cells were re-fixed with 4% formaldehyde in PBS for 15 minutes. Coverslips were then washed with PBS two times and incubated with RNase A (Qiagen, Cat# 19101) in PBS at 37 °C for one hour. After RNA removal, coverslips were dehydrated by successively incubating in 70%, 85%, 95%, and 100% ethanol for one minute each. Dried coverslips were incubated with FISH hybridization buffer (50% formamide (v/v), 2X SSC, 10% dextran sulfate (v/v), 100 ng/µL fragmented salmon sperm DNA, 5 pmol target probe). After DNA denaturation by incubating coverslips at 78 °C for 10 minutes in a thermocycler, coverslips were then incubated in the humidified chamber at 42 °C overnight. Next day, coverslips were washed four times with 2X SSC. The second wash was done with 2X SSC with 1 ng/µL HOECHST 33342 (Life Technologies, Cat# H3570). After the final wash, the coverslips were mounted on slides with mounting medium (Vector Laboratories, Cat# H-1000). Mounted coverslips were imaged with Leica TCS SP5 confocal microscope with at least two times line averaging for 512×512 pixels.

### Quantification of germ cells in histological sections

Seminiferous tubule stages were determined based on characteristic cellular organizations (Ahmed and de Rooij, 2009) after Periodic acid-Schiff (PAS), Hematoxylin staining of Bouin-fixed testis sections. A (A1 and undifferentiated) spermatogonia and pre-leptotene spermatocytes were counted in stage VII/VIII tubules. A (A3 and undifferentiated) spermatogonia and zygotene spermatocytes were counted in stage XII tubules. The size of the tubule perimeter was measured to normalize cell numbers to a standard tubule size. Only the sections cut perpendicular to the tubule axis were counted to have a consistent number of germ cells at the basement membrane for a given tubule perimeter. Cell numbers from at least 15 stage VII/VIII, and 10 stage XII tubules were counted per animal from two regions of a testis.

### Single cell RNA-seq

We used Chromium Single Cell 3’ Library & Gel Bead Kit v2 (10X Genomics, Cat: 120267) for single cell RNA-seq from wild-type adult testes. Testes were dissociated as described in the ‘*Testis dissociation to obtain single cells*’ section. Dissociated cells were diluted to 1000 cells/µL, and total 5200 cells (5.2 µL) were loaded into the Chromium controller to obtain RNA-seq profiles of approximately 3000 cells. After producing emulsions encapsulating gel beads and cells, reverse transcription and 12 cycles of PCR were performed to produce cell-barcoded amplified cDNA. Illumina sequencing-ready libraries were produced from 200 ng of the amplified cDNA based on the manufacturer’s protocol. 12 cycles of library amplification indexing PCR was performed.

### Genotyping of Cbx2 amplicon from single cell RNA-seq

Single cell analysis with targeted genotyping was done by adapting ’Genotyping of Transcriptome (GoT)’ (Nam et al., 2019) and V(D)J region amplification of T or B-cell receptor protocol by the 10X Genomics. We aimed to make specific *Cbx2* amplicons from cell-barcoded cDNA to amplify cDNA fragments and sequence Exon5 junctions to distinguish wild type (Exon4 spliced to Exon5) and mutant (Exon2 spliced to Exon5 after the deletion of Exon3, 4) for specific cell barcodes. c-KIT(+) spermatogonia were sorted to concentration of about 300∼600 cells/µL, and a total ∼8000 cells were loaded into the Chromium controller. Single cell barcoded cDNA was made with Chromium Next GEM Single Cell 5’ Reagent Kits (10X Genomics, Cat#:1000265) based on the manufacturer’s protocol with 14 PCR cycles of cDNA amplification. We used 5’ barcoding instead of 3’ barcoding because deleted exons of *Cbx2* were close to the 5’ end of the gene. Illumina sequencing-ready libraries were produced from 50 ng of the amplified cDNA based on the manufacturer’s protocol. 14 cycles of library amplification indexing PCR was performed.

To generate *Cbx2* amplicons, we performed nested PCRs to amplify *Cbx2*-specific cDNAs with cell barcodes. In the second PCR step of the nested PCR, *Cbx2*-specific primer that has a partial Illumina read2 sequence was used. The *Cbx2*-specific second nested primer also contains ’stagger’ sequences between the *Cbx2*-specific region and partial read2 to increase library complexity for accurate base calls during Illumina sequencing. We confirmed the amplification of the correct-sized bands corresponding to wild-type and deleted *Cbx2* amplicons by agarose gel electrophoresis. The amplified bands were excised and gel purified. Index PCR was performed to make the final Illumina-compatible libraries with unique indices with 10X provided index-PCR primers. *Cbx2* amplicon libraries were pooled with standard 10X single cell gene expression libraries to occupy ∼2% of total reads. A total of 4 samples were sequenced in one P3 lane on a NextSeq 2000 (NextSeq 2000 P3 Reagents (100 Cycles) Cat:20040559).

### CUT&RUN

CUT&RUN (ref)was performed with the following modifications. FACS purified, >100,000 c-KIT(+) spermatogonia were attached to Poly-D-Lysine (R&D Systems, Cat# 3439-200-01) coated 24-well plates by 1 minute centrifugation at 80 × g (instead of binding to Concanavalin A beads). After the attachment, cells were fixed with 1% formaldehyde in PBS for 15 minutes. After two PBS washes and one PBST (0.05% Triton X-100 (v/v)) wash, cells were incubated with primary antibodies in PBST BSA (0.05% Triton X-100, 3% BSA) overnight at 4 °C. Next day, cells were washed twice with PBST and finally washed with PBST EDTA (2 mM). Cells were then incubated with 500 ng/mL pA-MNase (protein A-MNase) in PBST BSA EDTA for more than 1 hour at 4 °C. After pA-MNase binding, cells were washed with PBST EDTA twice, and the plate was placed on wet ice. Cells were then incubated in pre-chilled 200 µL PBST with 2 mM CaCl_2_ for 30 minutes. After MNase cutting, 200 µL stop solution (340 mM NaCl, 20 mM EDTA, 4 mM EGTA, 50 µg/µL RNase A) was added, and the plate was placed in a 37 °C oven for 15 minutes. Released cut fragments were de-crosslinked in 0.1% SDS and 125 µg/mL Proteinase K (Sigma-Aldrich, Cat# 03115828001) in a 65 °C oven for >12 hours. DNA was phenol-chloroform extracted and sodium acetate/ethanol precipitated. Resuspended DNA was further cleaned up using 2.0X Solid Phase Reversible Immobilization (SPRI) beads to remove residual salts. Illumina sequencing libraries were generated based on a published protocol (Bowman et al., 2013) with the following modifications. SPRI beads were used in 2.0X ratio following end-repair and A-tailings steps instead of 1.8X. Adapters used in 2 nM final concentration instead of 10 nM. PCR extension was done for 30 instead of 45 seconds. Final PCR cycles were between 12 to 17. Number of PCR cycles were determined to be in the exponential phase by a pre-run of 10% of reaction with SYBR green dye in a real time PCR machine.

### Analysis of standard single cell RNA-seq

Adult testis (Figure 1): The raw single cell sequencing data generated for this paper are available at NCBI Geo (). Cellranger count (cellranger/2.0.0) was used to map fastq files to the mouse genome (refdata-cellranger=mm10-1.2.0). 141 million reads were processed to obtain 875 cells with 162K mean reads and 2593 median number of detected genes per cell. Ten principal components were used for t-SNE visualization and k-means clustering. Data were visualized by the Loupe browser (10X Genomics).

P15 testis (Figure 2): P15 single cell RNA-seq data were obtained from ArrayExpress (E-MTAB-6946, P15: do18195) (Ernst *et al*., 2019). Cellranger count (cellranger/3.0.2) was used to map fastq files to the mouse genome (refdata-cellranger-mm10-3.0.0) and obtain mtx files with UMI number-filtered raw expression values in each cell.

R package Seurat (R 3.6.3, Seurat 3.1.4) was used for all subsequent analyses (Stuart et al., 2019). Cells with more than 1000 detected RNA were used (nFeature_RNA > 1000) as previously done (Ernst *et al*., 2019). Ten principal components were used for clustering. From the initial clusters, spermatogonia and early spermatocytes were identified and chosen as a subset (3830 cells). The subset was re-clustered with 10 principal components and resolution 0.25. Dot plots for *Cbx2* and *Foxc2* expression level were drawn with Matlab (R2016a) after exporting data from R. All other plots were generated using Seurat. Default parameters were used for all above analyses unless otherwise specified.

### Analysis of amplicon single cell RNA-seq of Cbx2 inducible KO

Cellranger count (cellranger/6.0) was used to map fastq files to the mouse genome (refdata-cellranger-mm10-3.0.0) and to obtain gene expression counts per cell. R package Seurat (R 4.1.0, Seurat 4.0.4) was used for subsequent analyses (Hao et al., 2021). Cells were filtered by a) nFeature_RNA > 2000, b) mitochondrial RNA percentage < 7.5, c) percent occupied by the largest gene < 3, and d) RNA count and feature ratio < 5.5 (see scripts posted on the Github). Two control and two inducible KO samples were integrated to align cell populations across all four samples. Integrated data were then scaled and processed with 30 principal components for clustering. Differential gene expression tests were performed by the FindMarkers function of Seurat using Wilcoxon rank sum test.

*Cbx2* genotypes were assigned based on independent *Cbx2* amplicon sequencing data using a custom Python script. The script identifies ’**<u>TGGCGCATCTGGTTC</u>**CTTGAGCTTGGAGCG’ (WT, **<u>exon5</u>**linked to exon4) or ’**<u>TGGCGCATCTGGTTC</u>**TTGGAGGACCAGCCG’ (Mut, **<u>exon5</u>**linked to exon2) from the read2 of the paired-end reads and assigns the genotypes to the associated read1 (UMI-barcode). When an UMI is assigned to two different genotypes, likely due to PCR chimera formation, the genotype was left unassigned unless reads supporting one genotype are present 10 times more than the other genotype. A cell was assigned to be ’Mut’ when all *Cbx2* amplicon reads show recombination (exon5 linked to exon2), and otherwise assigned to be ’WTorHet’. Only the cells that have at least four UMIs linked to *Cbx2* sequence were assigned genotypes. The metadata for their genotypes (WT/Het or Mut) were added to the Seurat object by the AddMetaData function.

### Analysis of CUT&RUN data

The CUT&RUN sequencing data generated for this paper are available at NCBI GEO (). Paired-end fastq reads were quality trimmed (score 20), with stringency 1 using trim_galore (ver. 0.4.3) (Martin, 2011). Trimmed reads were mapped to the mm10 genome using bowtie2 (ver. 2.3.1) with the end-to-end option (Langmead and Salzberg, 2012). Mapped reads were qualify-filtered with MAPQ higher than 10, and only properly-paired reads were retained using Samtools (ver. 1.4.1) (Li et al., 2009). Picard (ver. 2.17.1) was used to remove duplicate reads (http://broadinstitute.github.io/picard). Alignment files (bam) were converted to bigwig files using Deeptools (ver. 2.2.4) by normalizing with RPKM (Ramirez et al., 2016). Peaks were identified by using MACS2 (ver. 2.1) with the ‘--broad’ option, by comparing CUT&RUN results using CBX2 antibodies (Experimental) to IgG (Control) or using HA antibodies applied to *Cbx2^HA/HA^* (Experimental) or *Cbx2^+/+^* (Control) mice (Zhang et al., 2008). CBX2 target genes were selected by first choosing the genes with transcription start sites overlapping with the CBX2 peaks and by only retaining the ones with more than one CBX2 (antibodies: 855, 856) or CBX2-HA (antibodies: 16B12, ab9110) CUT&RUN result showing that the gene is a target. Enrichment heat maps and numeric data for averaged plots were obtained using the computeMatrix tool from Deeptools. Averaged plots were drawn by Matlab. Promoter associated enrichment scores were obtained using the multiBigwigSummary function of deeptools.

All analysis scripts are available at (github.com/jongminkmg/cbx2).

## Supporting information

Key resources table

## Acknowledgements

We thank Minx Fuller and Fuller, Page and Kingston laboratory members for insightful discussions. We thank Dirk de Rooij and Tsutomu Endo for advice on mammalian spermatogenesis, Lin Wu (Harvard Genome Modification Facility) for *HA-Cbx2* knock-in mice generation, Manashree Damle for advice on bioinformatic analyses, Jen Sheen, Susan Cotman and Jeannie Lee laboratories for help and access to microscopes. We thank MacKenzie Mauger, Wojciech Siwek, Ukrae Cho and Julia Wucherpfennig for careful reading of the manuscript. J.J.K. was supported by the Urology Care Foundation Research Scholar Award Program 2018A012748, M.S.L. was supported by the Agency of Science, Research and Technology, Singapore, and D.C.P. was supported by Howard Hughes Medical Institute. This work was supported by the NIH grants R01GM4390121A1 and R35GM131743 to R.E.K. The content is the responsibility of the authors and does not represent the views of the funders.

## Author contributions

J.J.K. and R.E.K conceived the study. J.J.K. and E.R.S. performed experiments and data analysis. M.S.L. generated *Cbx2^Δ^* mice. D.C.P. advised on characterization of *Cbx2* mutant testis phenotypes. J.J.K. and R.E.K. wrote the manuscript with input from all authors.

---

**Figure S1.**
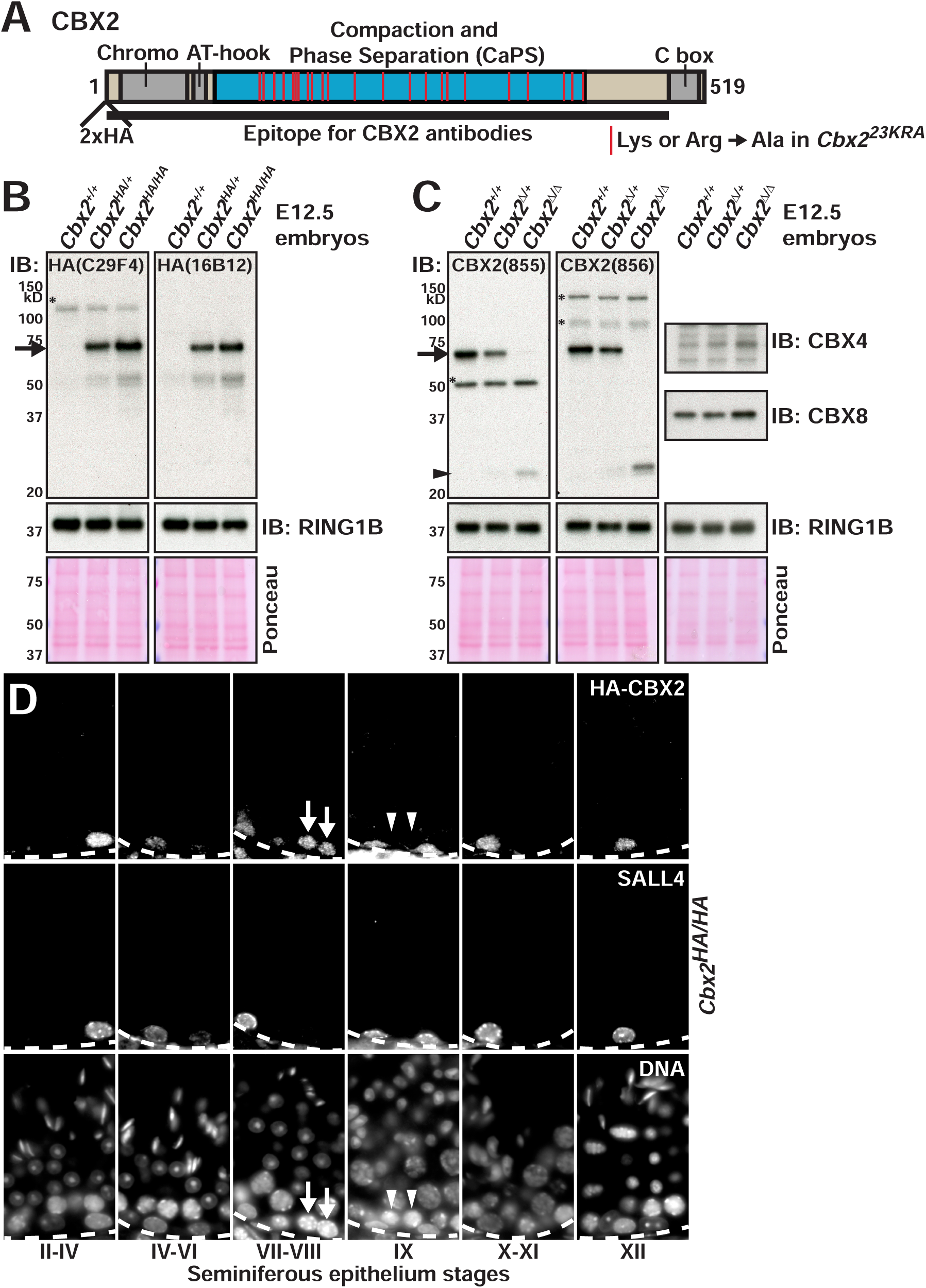
Specificity of CBX2 and HA antibodies against endogenous and HA-tagged CBX2 (A) Schematic domain structure of CBX2. The positions of Lys and Arg replaced with Ala in the *Cbx2^23KRA^* mutant are marked by red bars. (B and C) Western blot of CBX2 using (B) HA antibodies with wild-type, *Cbx2^HA/+^*, *Cbx2^HA/HA^* and (C) affinity purified CBX2 antibodies with wild-type, *Cbx2^Δ/+^*, *Cbx2^Δ/Δ^* embryonic day 12.5 (E12.5) embryo lysates. The HA knock-in mouse (*Cbx2^HA/HA^*) was homozygous-viable and fertile. RING1B was blotted to show the total PRC1 complex amount. Ponceau staining was used as the loading control. (C) A full-length CBX2 band was detected at right below the 75 kD size marker (arrow), and a truncated CBX2 band was detected at slightly above the 20 kD size marker from *Cbx2* deletion embryos (arrowhead). CBX4 and CBX8 were blotted to show potential paralogous compensation in *Cbx2* deletion embryos. Asterisks (*) denote non-specific background bands. (D) Co-immunofluorescence staining of testis sections of a *Cbx2^HA/HA^*animal with HA (C29F4) and SALL4 antibodies. Different seminiferous tubule stages were denoted at the bottom. Dotted lines represent the basement membrane. Arrows indicate pre-leptotene spermatocytes (CBX2+) and arrowheads indicate leptotene spermatocytes (CBX2-).

**Figure S2.**
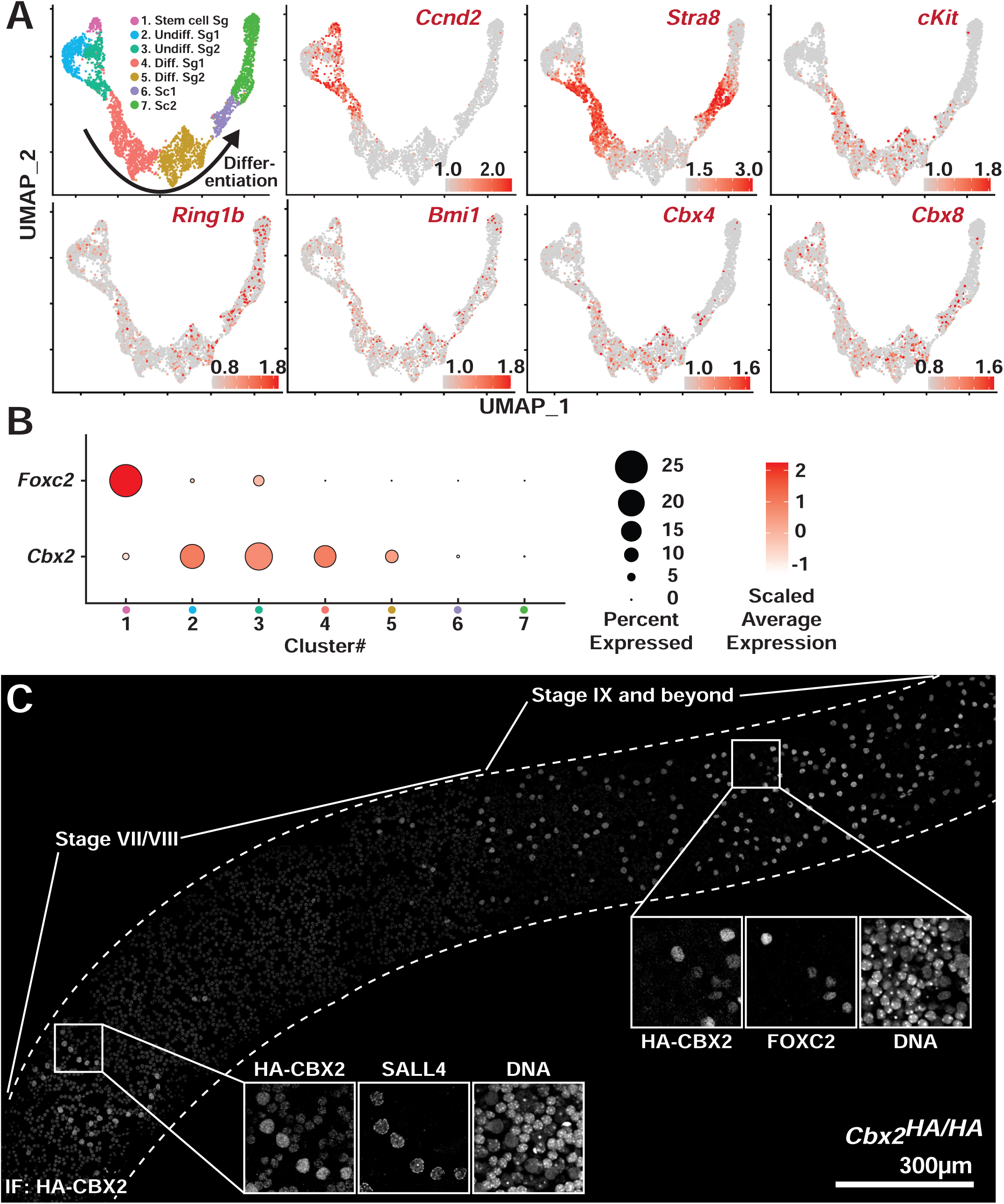
CBX2 is upregulated as spermatogonial stem cells differentiate (A) UMAP representation of a subset of spermatogonia and early spermatocytes from scRNA-seq from p15 mouse testes shown in Fig. 2B (Ernst *et al*., 2019). Normalized expression levels of *Ccnd2* (enriched in a subset of undifferentiated spermatogonia and A1 spermatogonia), *Stra8* (upregulated first in A1 spermatogonia and later in pre-leptotene spermatocytes), *c-Kit* (enriched in differentiating spermatogonia), and *Ring1b*/*Bmi1*/*Cbx4/Cbx8* (components of cPRC1) are represented in red. (B) Dot plots showing the percentage (dot size) of cells expressing *Foxc2* or *Cbx2* in each cluster and scaled average expression (shade of red). Data are scaled to have mean 0 with standard deviation 1. (C) Co-immunofluorescence staining of a whole testis tubule of *Cbx2^HA/HA^*animals with HA, SALL4, and FOXC2 antibodies. CBX2 signal was represented in contiguous tubule stages from VII to IX and beyond. The inset from stage VII/VIII is an example of SALL4(+) A1 spermatogonia, and smaller pre-leptotene spermatocytes with CBX2 signal. The inset from stage IX and beyond is an example of As FOXC2(+) spermatogonia without CBX2 signal and Aaln4 FOXC2(+) spermatogonia with weak CBX2 signal.

**Figure S3.**
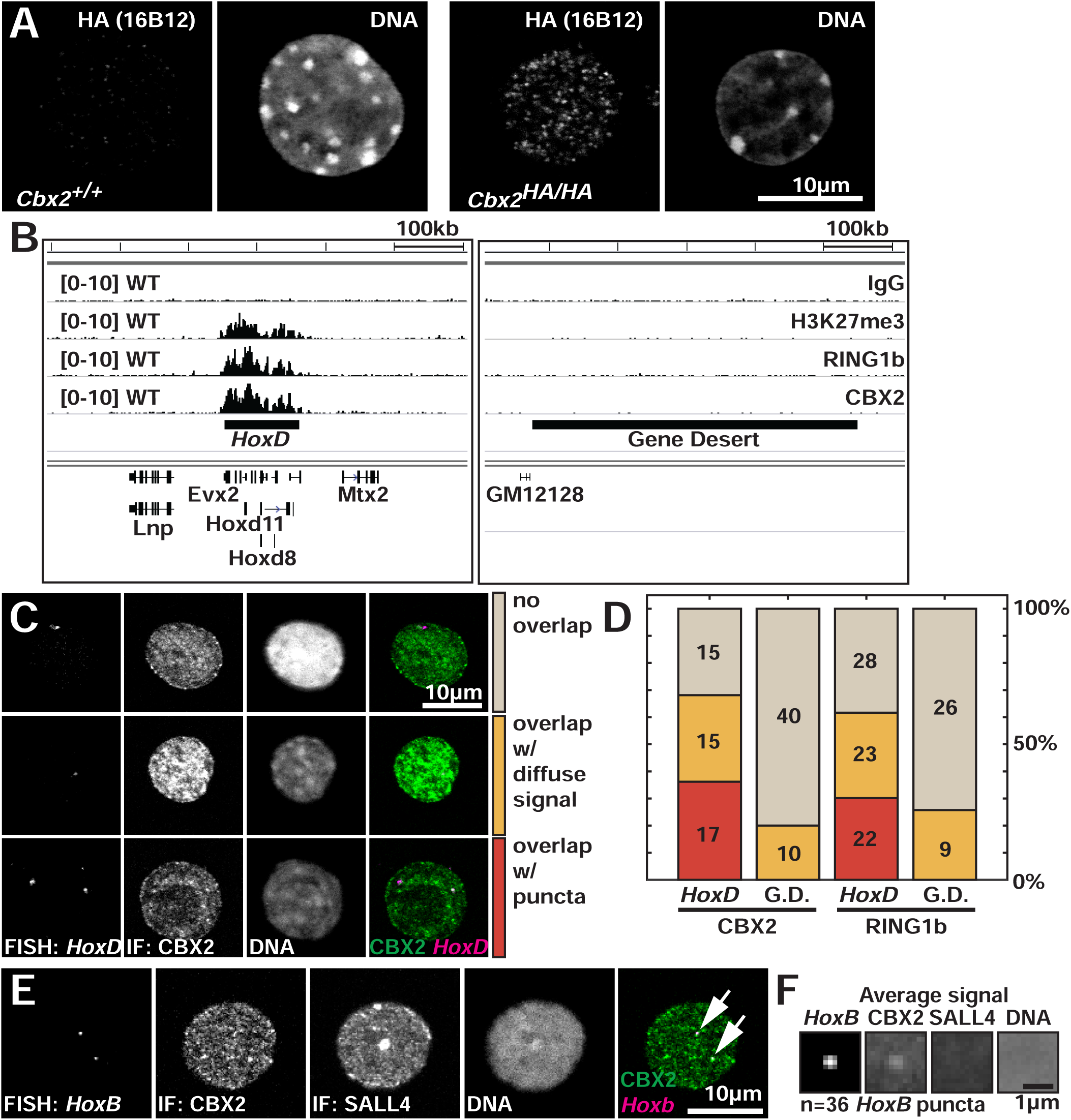
*HoxD* loci overlap with CBX2 puncta (A) Immunostaining of HA epitope using FACS sorted c-KIT(+) spermatogonia from *Cbx2^+/+^* and *Cbx2^HA/HA^* mice. (B) Genome browser screenshots of CUT&RUN enrichment of IgG, H3K27me3, RING1B, and CBX2 in FACS sorted c-KIT(+) spermatogonia at *HoxD* and gene desert loci. (C) Examples with different patterns of overlap between *HoxD* locus and CBX2 by co-immuno-FISH. (D) Quantification of different overlap patterns between *HoxD* or gene desert loci with CBX2 or RING1B. (E) Co-immuno-FISH of *HoxB* locus and CBX2 and SALL4 protein using FACS sorted c-KIT(+) spermatogonia from wild-type mice. A single optical section was represented. Arrows represent *HoxB* loci overlapping with CBX2 puncta. (F) Average signal projection of 2 µm square centered at *HoxB* puncta. Average signal projection of CBX2, SALL4, and DNA at the corresponding locations were also represented.

**Figure S4.**
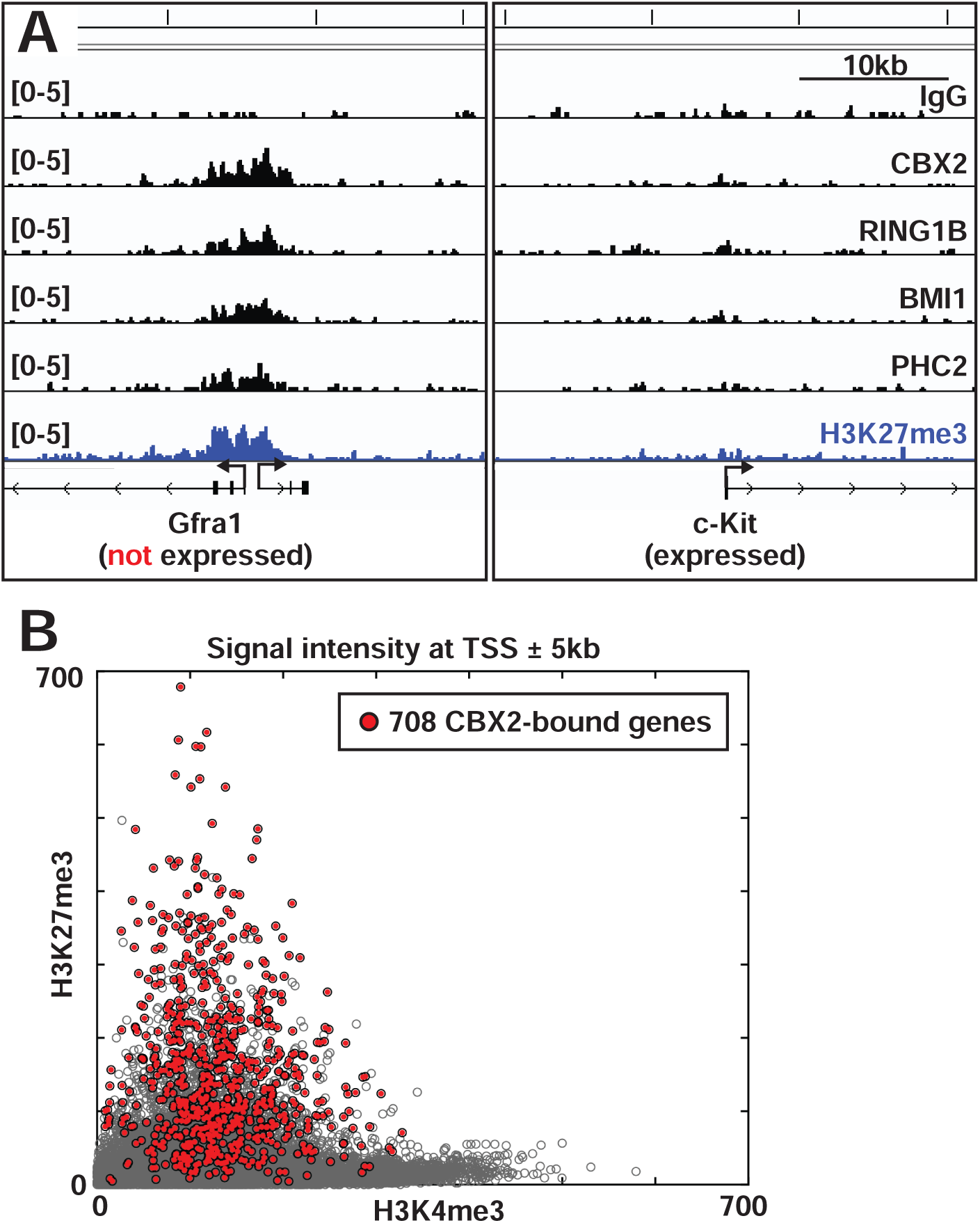
CBX2 and H3K27me3 are depleted at genes with strong H4K3me3 enrichment (A) Additional example genome browser screenshots of CUT&RUN enrichment of IgG, CBX2, cPRC1 components (RING1B, BMI1, PHC2), and H3K27me3 from FACS sorted c-KIT(+) spermatogonia. Enrichment profiles at *Gfra1* and *c-Kit* genes are shown as representative examples. *Gfra1* is highly expressed in spermatogonial stem cells and downregulated in c-KIT(+) differentiating spermatogonia. In contrast, *c-Kit* is robustly expressed in c-KIT(+) spermatogonia. (B) A scatter plot showing intensity relationship between H3K4me3 and H3K27me3 at 10kb regions encompassing 28,656 transcription start sites (open gray circles). CBX2-bound genes were represented with red-filled circles.

**Figure S5.**
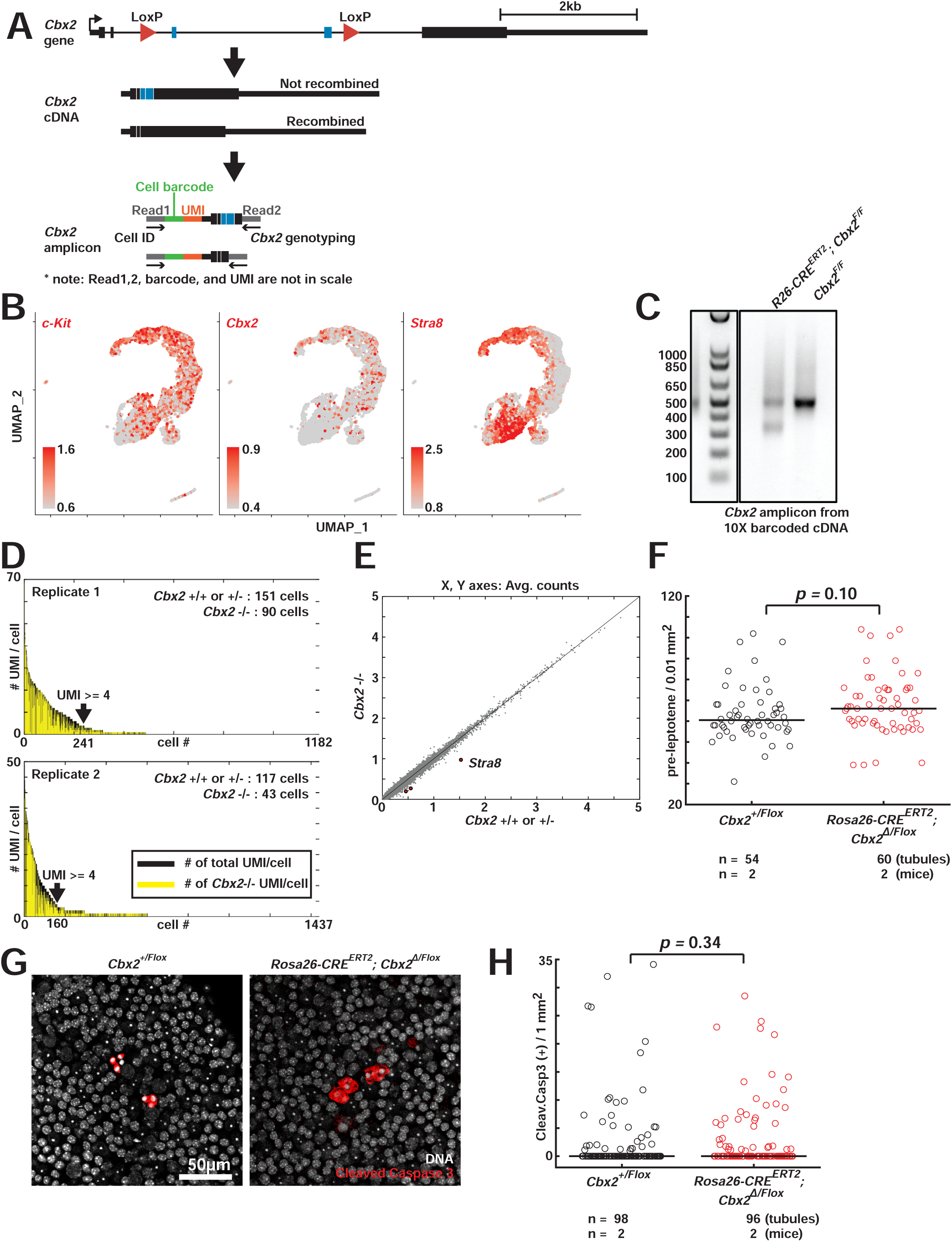
CBX2 is required for differentiation to CCND2(+) A1 spermatogonia (A) Schematic of genotyping strategy by sequencing *Cbx2* cDNA amplicons with cell barcodes. In the *Cbx2* gene schematic, thick lines represent five *Cbx2* exons (based on NM_007623.3), and thin lines represent introns. Blue lines represent exon3 and exon4 that are deleted when recombined by CRE. *Cbx2* amplicons were amplified using *Cbx2*-specific primers such that the final product included the *Cbx2* exon5 junctions and the cell barcodes. (B) UMAP representation showing normalized expression levels of *c-Kit*, *Cbx2*, and *Stra8* in FACS sorted c-KIT(+) spermatogonia. The UMAPs represent combined cell profiles from control and *Cbx2* inducible mutant animals. (C) A DNA gel image of *Cbx2* amplicons generated from 10X cell-barcoded cDNAs from control and *Cbx2* inducible mutant animals. (D) Distribution of the number of total UMIs detected per cell (black lines) and the number of UMIs associated with *Cbx2* mutation (yellow lines). Data are sorted by the decreasing amount of total UMIs per cell from left to right. Cellular genotypes are assigned only when ≥4 UMIs are detected per cell. (E) A scatter plot comparing averaged gene expression levels of *Cbx2*-expressing cells (population#1, #2, and #3 in Fig. 5B) between *Cbx2^+/+, +/-^*and *Cbx2^-/-^* cells. Red dots represent three differentially expressed genes (*p* < 0.05, after Bonferroni correction) between 268 *Cbx2^+/+, +/-^* cells versus 133 *Cbx2^-/-^* cells (Wilcoxon rank sum test). (F) Quantification of the number of pre-leptotene spermatocytes per 0.01mm^2^ segments of stage VII/VIII tubule sections. *p* value was calculated by two-tailed t-test. (G) Immunofluorescence staining of whole testis tubules of control (*Cbx2^+/Flox^*) and *Cbx2* inducible mutant (*Rosa26-CRE^ERT2^; Cbx2^Δ/Flox^*) animals with cleaved Caspase 3 (red) antibody to mark apoptotic cells. Only stage VI to VIII tubules (where A aligned to A1 transition happens) are analyzed. (H) Quantification of the number of apoptotic cells with cleaved Caspase 3 signal per 1mm^2^ segments of stage VI to VIII tubule sections. *p* value was calculated by Wilcoxon rank sum test.

**Figure S6.**
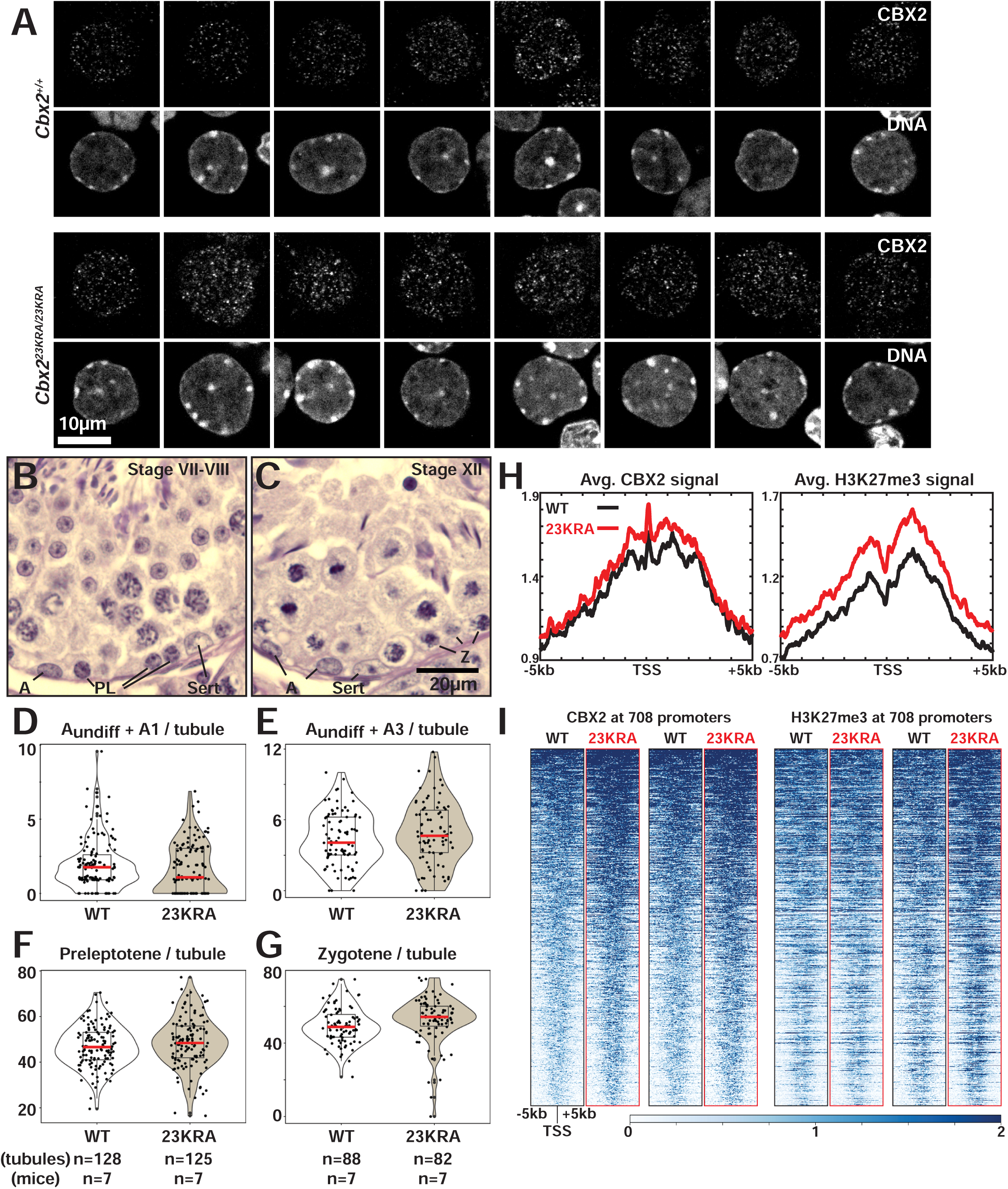
No significant differences in nuclear and chromatin distribution of CBX2 between wild-type and *Cbx2* CAPS domain mutant mice (A) Immunostaining of CBX2 using FACS sorted c-KIT(+) spermatogonia from wild-type and *Cbx2^23KRA/23KRA^* mice. Spermatogonia at a similar stage between wild-type and *Cbx2^23KRA/23KRA^*were chosen based on nuclear morphology. Eight representative nuclei per condition were shown. (B and C) Representative Hematoxylin-PAS staining showing part of tubule sections at (B) stage VII/VIII and (C) stage XII. A (at stage VII/VIII): undifferentiated and A1 spermatogonia; A (at stage XII): undifferentiated and A3 spermatogonia; PL: pre-leptotene spermatocytes; Z: zygotene spermatocytes; Sert: Sertoli cells. (D-G) Quantification of the number of (D) undifferentiated and A1 spermatogonia, (E) undifferentiated and A3 spermatogonia, (F) pre-leptotene spermatocytes, (G) zygotene spermatocytes in (D and F) stage VII/VIII or (E and G) XII tubules in wild-type and *Cbx2^23KRA/23KRA^* mice after normalizing cell numbers to average tubule circumference. (H) Average CUT&RUN enrichment of CBX2 and H3K27me3 using FACS-sorted c-KIT(+) spermatogonia. Signals centered at promoters of 708 CBX2 target promoters in wild-type (black lines) and *Cbx2^23KRA/23KRA^*mutant (red lines) spermatogonia are represented. (I) Heatmap of profiles at individual 708 genes of (H).

## References

Aloisio, G.M., Nakada, Y., Saatcioglu, H.D., Pena, C.G., Baker, M.D., Tarnawa, E.D., Mukherjee, J., Manjunath, H., Bugde, A., Sengupta, A.L., et al. (2014). PAX7 expression defines germline stem cells in the adult testis. J Clin Invest 124, 3929–3944. 10.1172/JCI75943.

Bantignies, F., Roure, V., Comet, I., Leblanc, B., Schuettengruber, B., Bonnet, J., Tixier, V., Mas, A., and Cavalli, G. (2011). Polycomb-dependent regulatory contacts between distant Hox loci in Drosophila. Cell 144, 214–226. 10.1016/j.cell.2010.12.026.

Beliveau, B.J., Joyce, E.F., Apostolopoulos, N., Yilmaz, F., Fonseka, C.Y., McCole, R.B., Chang, Y., Li, J.B., Senaratne, T.N., Williams, B.R., et al. (2012). Versatile design and synthesis platform for visualizing genomes with Oligopaint FISH probes. Proc Natl Acad Sci U S A 109, 21301–21306. 10.1073/pnas.1213818110.

Beumer, T.L., Roepers-Gajadien, H.L., Gademan, I.S., Kal, H.B., and de Rooij, D.G. (2000). Involvement of the D-type cyclins in germ cell proliferation and differentiation in the mouse. Biol Reprod 63, 1893–1898. 10.1095/biolreprod63.6.1893.

Blackledge, N.P., and Klose, R.J. (2021). The molecular principles of gene regulation by Polycomb repressive complexes. Nat Rev Mol Cell Biol 22, 815–833. 10.1038/s41580-021-00398-y.

Bowman, S.K., Simon, M.D., Deaton, A.M., Tolstorukov, M., Borowsky, M.L., and Kingston, R.E. (2013). Multiplexed Illumina sequencing libraries from picogram quantities of DNA. BMC Genomics 14, 466. 10.1186/1471-2164-14-466.

Boyer, L.A., Plath, K., Zeitlinger, J., Brambrink, T., Medeiros, L.A., Lee, T.I., Levine, S.S., Wernig, M., Tajonar, A., Ray, M.K., et al. (2006). Polycomb complexes repress developmental regulators in murine embryonic stem cells. Nature 441, 349–353. 10.1038/nature04733.

Buaas, F.W., Kirsh, A.L., Sharma, M., McLean, D.J., Morris, J.L., Griswold, M.D., de Rooij, D.G., and Braun, R.E. (2004). Plzf is required in adult male germ cells for stem cell self-renewal. Nat Genet 36, 647–652. 10.1038/ng1366.

Cao, R., Wang, L., Wang, H., Xia, L., Erdjument-Bromage, H., Tempst, P., Jones, R.S., and Zhang, Y. (2002). Role of histone H3 lysine 27 methylation in Polycomb-group silencing. Science 298, 1039–1043. 10.1126/science.1076997.

Chiacchiera, F., Rossi, A., Jammula, S., Piunti, A., Scelfo, A., Ordonez-Moran, P., Huelsken, J., Koseki, H., and Pasini, D. (2016). Polycomb Complex PRC1 Preserves Intestinal Stem Cell Identity by Sustaining Wnt/beta-Catenin Transcriptional Activity. Cell Stem Cell 18, 91–103. 10.1016/j.stem.2015.09.019.

Core, N., Bel, S., Gaunt, S.J., Aurrand-Lions, M., Pearce, J., Fisher, A., and Djabali, M. (1997). Altered cellular proliferation and mesoderm patterning in Polycomb-M33-deficient mice. Development 124, 721–729.

Costoya, J.A., Hobbs, R.M., Barna, M., Cattoretti, G., Manova, K., Sukhwani, M., Orwig, K.E., Wolgemuth, D.J., and Pandolfi, P.P. (2004). Essential role of Plzf in maintenance of spermatogonial stem cells. Nat Genet 36, 653–659. 10.1038/ng1367.

Czermin, B., Melfi, R., McCabe, D., Seitz, V., Imhof, A., and Pirrotta, V. (2002). Drosophila enhancer of Zeste/ESC complexes have a histone H3 methyltransferase activity that marks chromosomal Polycomb sites. Cell 111, 185–196. 10.1016/s0092-8674(02)00975-3.

Dauber, K.L., Perdigoto, C.N., Valdes, V.J., Santoriello, F.J., Cohen, I., and Ezhkova, E. (2016). Dissecting the Roles of Polycomb Repressive Complex 2 Subunits in the Control of Skin Development. J Invest Dermatol 136, 1647–1655. 10.1016/j.jid.2016.02.809.

de Napoles, M., Mermoud, J.E., Wakao, R., Tang, Y.A., Endoh, M., Appanah, R., Nesterova, T.B., Silva, J., Otte, A.P., Vidal, M., et al. (2004). Polycomb group proteins Ring1A/B link ubiquitylation of histone H2A to heritable gene silencing and X inactivation. Dev Cell 7, 663–676. 10.1016/j.devcel.2004.10.005.

de Rooij, D.G., and Russell, L.D. (2000). All you wanted to know about spermatogonia but were afraid to ask. J Androl 21, 776–798.

Endo, T., Romer, K.A., Anderson, E.L., Baltus, A.E., de Rooij, D.G., and Page, D.C. (2015). Periodic retinoic acid-STRA8 signaling intersects with periodic germ-cell competencies to regulate spermatogenesis. Proc Natl Acad Sci U S A 112, E2347–2356. 10.1073/pnas.1505683112.

Endoh, M., Endo, T.A., Endoh, T., Fujimura, Y., Ohara, O., Toyoda, T., Otte, A.P., Okano, M., Brockdorff, N., Vidal, M., and Koseki, H. (2008). Polycomb group proteins Ring1A/B are functionally linked to the core transcriptional regulatory circuitry to maintain ES cell identity. Development 135, 1513–1524. 10.1242/dev.014340.

Ernst, C., Eling, N., Martinez-Jimenez, C.P., Marioni, J.C., and Odom, D.T. (2019). Staged developmental mapping and X chromosome transcriptional dynamics during mouse spermatogenesis. Nat Commun 10, 1251. 10.1038/s41467-019-09182-1.

Fischle, W., Wang, Y., Jacobs, S.A., Kim, Y., Allis, C.D., and Khorasanizadeh, S. (2003). Molecular basis for the discrimination of repressive methyl-lysine marks in histone H3 by Polycomb and HP1 chromodomains. Genes Dev 17, 1870–1881. 10.1101/gad.1110503.

Flora, P., Dalal, G., Cohen, I., and Ezhkova, E. (2021). Polycomb Repressive Complex(es) and Their Role in Adult Stem Cells. Genes (Basel) 12. 10.3390/genes12101485.

Francis, N.J., Kingston, R.E., and Woodcock, C.L. (2004). Chromatin compaction by a polycomb group protein complex. Science 306, 1574–1577. 10.1126/science.1100576.

Gao, Z., Zhang, J., Bonasio, R., Strino, F., Sawai, A., Parisi, F., Kluger, Y., and Reinberg, D. (2012). PCGF homologs, CBX proteins, and RYBP define functionally distinct PRC1 family complexes. Mol Cell 45, 344–356. 10.1016/j.molcel.2012.01.002.

Garcia-Moreno, S.A., Lin, Y.T., Futtner, C.R., Salamone, I.M., Capel, B., and Maatouk, D.M. (2019). CBX2 is required to stabilize the testis pathway by repressing Wnt signaling. PLoS Genet 15, e1007895. 10.1371/journal.pgen.1007895.

Gely-Pernot, A., Raverdeau, M., Celebi, C., Dennefeld, C., Feret, B., Klopfenstein, M., Yoshida, S., Ghyselinck, N.B., and Mark, M. (2012). Spermatogonia differentiation requires retinoic acid receptor gamma. Endocrinology 153, 438–449. 10.1210/en.2011-1102.

Gendall, A.R., Levy, Y.Y., Wilson, A., and Dean, C. (2001). The VERNALIZATION 2 gene mediates the epigenetic regulation of vernalization in Arabidopsis. Cell 107, 525–535. 10.1016/s0092-8674(01)00573-6.

Grau, D.J., Chapman, B.A., Garlick, J.D., Borowsky, M., Francis, N.J., and Kingston, R.E. (2011). Compaction of chromatin by diverse Polycomb group proteins requires localized regions of high charge. Genes Dev 25, 2210–2221. 10.1101/gad.17288211.

Grimaud, C., Bantignies, F., Pal-Bhadra, M., Ghana, P., Bhadra, U., and Cavalli, G. (2006). RNAi components are required for nuclear clustering of Polycomb group response elements. Cell 124, 957–971. 10.1016/j.cell.2006.01.036.

Hao, Y., Hao, S., Andersen-Nissen, E., Mauck, W.M., 3rd, Zheng, S., Butler, A., Lee, M.J., Wilk, A.J., Darby, C., Zager, M., et al. (2021). Integrated analysis of multimodal single-cell data. Cell 184, 3573–3587 e3529. 10.1016/j.cell.2021.04.048.

Hirabayashi, Y., Suzki, N., Tsuboi, M., Endo, T.A., Toyoda, T., Shinga, J., Koseki, H., Vidal, M., and Gotoh, Y. (2009). Polycomb limits the neurogenic competence of neural precursor cells to promote astrogenic fate transition. Neuron 63, 600–613. 10.1016/j.neuron.2009.08.021.

Hofmann, M.C., Braydich-Stolle, L., and Dym, M. (2005). Isolation of male germ-line stem cells; influence of GDNF. Dev Biol 279, 114–124. 10.1016/j.ydbio.2004.12.006.

Ikami, K., Tokue, M., Sugimoto, R., Noda, C., Kobayashi, S., Hara, K., and Yoshida, S. (2015). Hierarchical differentiation competence in response to retinoic acid ensures stem cell maintenance during mouse spermatogenesis. Development 142, 1582–1592. 10.1242/dev.118695.

Iovino, N., Ciabrelli, F., and Cavalli, G. (2013). PRC2 controls Drosophila oocyte cell fate by repressing cell cycle genes. Dev Cell 26, 431–439. 10.1016/j.devcel.2013.06.021.

Isono, K., Endo, T.A., Ku, M., Yamada, D., Suzuki, R., Sharif, J., Ishikura, T., Toyoda, T., Bernstein, B.E., and Koseki, H. (2013). SAM domain polymerization links subnuclear clustering of PRC1 to gene silencing. Dev Cell 26, 565–577. 10.1016/j.devcel.2013.08.016.

Jacobs, J.J., Scheijen, B., Voncken, J.W., Kieboom, K., Berns, A., and van Lohuizen, M. (1999). Bmi-1 collaborates with c-Myc in tumorigenesis by inhibiting c-Myc-induced apoptosis via INK4a/ARF. Genes Dev 13, 2678–2690. 10.1101/gad.13.20.2678.

Jaensch, E.S., Zhu, J., Cochrane, J.C., Marr, S.K., Oei, T.A., Damle, M., McCaslin, E.Z., and Kingston, R.E. (2021). A Polycomb domain found in committed cells impairs differentiation when introduced into PRC1 in pluripotent cells. Mol Cell 81, 4677–4691 e4678. 10.1016/j.molcel.2021.09.018.

Kanatsu-Shinohara, M., Ogonuki, N., Inoue, K., Miki, H., Ogura, A., Toyokuni, S., and Shinohara, T. (2003). Long-term proliferation in culture and germline transmission of mouse male germline stem cells. Biol Reprod 69, 612–616. 10.1095/biolreprod.103.017012.

Katoh-Fukui, Y., Baba, T., Sato, T., Otake, H., Nagakui-Noguchi, Y., Shindo, M., Suyama, M., Ohkawa, Y., Tsumura, H., Morohashi, K.I., and Fukami, M. (2019). Mouse polycomb group gene Cbx2 promotes osteoblastic but suppresses adipogenic differentiation in postnatal long bones. Bone 120, 219–231. 10.1016/j.bone.2018.10.021.

Katoh-Fukui, Y., Owaki, A., Toyama, Y., Kusaka, M., Shinohara, Y., Maekawa, M., Toshimori, K., and Morohashi, K. (2005). Mouse Polycomb M33 is required for splenic vascular and adrenal gland formation through regulating Ad4BP/SF1 expression. Blood 106, 1612–1620. 10.1182/blood-2004-08-3367.

Katoh-Fukui, Y., Tsuchiya, R., Shiroishi, T., Nakahara, Y., Hashimoto, N., Noguchi, K., and Higashinakagawa, T. (1998). Male-to-female sex reversal in M33 mutant mice. Nature 393, 688–692. 10.1038/31482.

Kim, J.J., and Kingston, R.E. (2022). Context-specific Polycomb mechanisms in development. Nat Rev Genet. 10.1038/s41576-022-00499-0.

Kishi, J.Y., Lapan, S.W., Beliveau, B.J., West, E.R., Zhu, A., Sasaki, H.M., Saka, S.K., Wang, Y., Cepko, C.L., and Yin, P. (2019). SABER amplifies FISH: enhanced multiplexed imaging of RNA and DNA in cells and tissues. Nat Methods 16, 533–544. 10.1038/s41592-019-0404-0.

Komai, Y., Tanaka, T., Tokuyama, Y., Yanai, H., Ohe, S., Omachi, T., Atsumi, N., Yoshida, N., Kumano, K., Hisha, H., et al. (2014). Bmi1 expression in long-term germ stem cells. Sci Rep 4, 6175. 10.1038/srep06175.

Koppens, M.A., Bounova, G., Gargiulo, G., Tanger, E., Janssen, H., Cornelissen-Steijger, P., Blom, M., Song, J.Y., Wessels, L.F., and van Lohuizen, M. (2016). Deletion of Polycomb Repressive Complex 2 From Mouse Intestine Causes Loss of Stem Cells. Gastroenterology 151, 684–697 e612. 10.1053/j.gastro.2016.06.020.

Kotaja, N., Kimmins, S., Brancorsini, S., Hentsch, D., Vonesch, J.L., Davidson, I., Parvinen, M., and Sassone-Corsi, P. (2004). Preparation, isolation and characterization of stage-specific spermatogenic cells for cellular and molecular analysis. Nat Methods 1, 249–254. 10.1038/nmeth1204-249.

Koubova, J., Menke, D.B., Zhou, Q., Capel, B., Griswold, M.D., and Page, D.C. (2006). Retinoic acid regulates sex-specific timing of meiotic initiation in mice. Proc Natl Acad Sci U S A 103, 2474–2479. 10.1073/pnas.0510813103.

Kundu, S., Ji, F., Sunwoo, H., Jain, G., Lee, J.T., Sadreyev, R.I., Dekker, J., and Kingston, R.E. (2017). Polycomb Repressive Complex 1 Generates Discrete Compacted Domains that Change during Differentiation. Mol Cell 65, 432–446 e435. 10.1016/j.molcel.2017.01.009.

Kuzmichev, A., Nishioka, K., Erdjument-Bromage, H., Tempst, P., and Reinberg, D. (2002). Histone methyltransferase activity associated with a human multiprotein complex containing the Enhancer of Zeste protein. Genes Dev 16, 2893–2905. 10.1101/gad.1035902.

Langmead, B., and Salzberg, S.L. (2012). Fast gapped-read alignment with Bowtie 2. Nat Methods 9, 357–359. 10.1038/nmeth.1923.

Larson, A.G., Elnatan, D., Keenen, M.M., Trnka, M.J., Johnston, J.B., Burlingame, A.L., Agard, D.A., Redding, S., and Narlikar, G.J. (2017). Liquid droplet formation by HP1alpha suggests a role for phase separation in heterochromatin. Nature 547, 236–240. 10.1038/nature22822.

Lau, M.S. (2016). Mutations in the Charged Domain of CBX2 Disrupt PRC1 Function in Vivo. Doctoral (Harvard University).

Lau, M.S., Schwartz, M.G., Kundu, S., Savol, A.J., Wang, P.I., Marr, S.K., Grau, D.J., Schorderet, P., Sadreyev, R.I., Tabin, C.J., and Kingston, R.E. (2017). Mutation of a nucleosome compaction region disrupts Polycomb-mediated axial patterning. Science 355, 1081–1084. 10.1126/science.aah5403.

Lessard, J., Baban, S., and Sauvageau, G. (1998). Stage-specific expression of polycomb group genes in human bone marrow cells. Blood 91, 1216–1224.

Lessard, J., and Sauvageau, G. (2003). Bmi-1 determines the proliferative capacity of normal and leukaemic stem cells. Nature 423, 255–260. 10.1038/nature01572.

Levine, S.S., Weiss, A., Erdjument-Bromage, H., Shao, Z., Tempst, P., and Kingston, R.E. (2002). The core of the polycomb repressive complex is compositionally and functionally conserved in flies and humans. Mol Cell Biol 22, 6070–6078. 10.1128/MCB.22.17.6070-6078.2002.

Li, H., Handsaker, B., Wysoker, A., Fennell, T., Ruan, J., Homer, N., Marth, G., Abecasis, G., Durbin, R., and Genome Project Data Processing, S. (2009). The Sequence Alignment/Map format and SAMtools. Bioinformatics 25, 2078–2079. 10.1093/bioinformatics/btp352.

Li, L., and Clevers, H. (2010). Coexistence of quiescent and active adult stem cells in mammals. Science 327, 542–545. 10.1126/science.1180794.

Lovasco, L.A., Gustafson, E.A., Seymour, K.A., de Rooij, D.G., and Freiman, R.N. (2015). TAF4b is required for mouse spermatogonial stem cell development. Stem Cells 33, 1267–1276. 10.1002/stem.1914.

Maezawa, S., Hasegawa, K., Yukawa, M., Sakashita, A., Alavattam, K.G., Andreassen, P.R., Vidal, M., Koseki, H., Barski, A., and Namekawa, S.H. (2017). Polycomb directs timely activation of germline genes in spermatogenesis. Genes Dev 31, 1693–1703. 10.1101/gad.302000.117.

Martin, M. (2011). Cutadapt removes adapter sequences from high-throughput sequencing reads. EMBnet.journal 17. 10.14806/ej.17.1.200.

Mieczkowski, J., Cook, A., Bowman, S.K., Mueller, B., Alver, B.H., Kundu, S., Deaton, A.M., Urban, J.A., Larschan, E., Park, P.J., et al. (2016). MNase titration reveals differences between nucleosome occupancy and chromatin accessibility. Nat Commun 7, 11485. 10.1038/ncomms11485.

Min, J., Zhang, Y., and Xu, R.M. (2003). Structural basis for specific binding of Polycomb chromodomain to histone H3 methylated at Lys 27. Genes Dev 17, 1823–1828. 10.1101/gad.269603.

Molofsky, A.V., Pardal, R., Iwashita, T., Park, I.K., Clarke, M.F., and Morrison, S.J. (2003). Bmi-1 dependence distinguishes neural stem cell self-renewal from progenitor proliferation. Nature 425, 962–967. 10.1038/nature02060.

Morey, L., Pascual, G., Cozzuto, L., Roma, G., Wutz, A., Benitah, S.A., and Di Croce, L. (2012). Nonoverlapping functions of the Polycomb group Cbx family of proteins in embryonic stem cells. Cell Stem Cell 10, 47–62. 10.1016/j.stem.2011.12.006.

Mu, W., Starmer, J., Fedoriw, A.M., Yee, D., and Magnuson, T. (2014). Repression of the soma-specific transcriptome by Polycomb-repressive complex 2 promotes male germ cell development. Genes Dev 28, 2056–2069. 10.1101/gad.246124.114.

Muller, J., Hart, C.M., Francis, N.J., Vargas, M.L., Sengupta, A., Wild, B., Miller, E.L., O’Connor, M.B., Kingston, R.E., and Simon, J.A. (2002). Histone methyltransferase activity of a Drosophila Polycomb group repressor complex. Cell 111, 197–208. 10.1016/s0092-8674(02)00976-5.

Nam, A.S., Kim, K.T., Chaligne, R., Izzo, F., Ang, C., Taylor, J., Myers, R.M., Abu-Zeinah, G., Brand, R., Omans, N.D., et al. (2019). Somatic mutations and cell identity linked by Genotyping of Transcriptomes. Nature 571, 355–360. 10.1038/s41586-019-1367-0.

O’Loghlen, A., Munoz-Cabello, A.M., Gaspar-Maia, A., Wu, H.A., Banito, A., Kunowska, N., Racek, T., Pemberton, H.N., Beolchi, P., Lavial, F., et al. (2012). MicroRNA regulation of Cbx7 mediates a switch of Polycomb orthologs during ESC differentiation. Cell Stem Cell 10, 33–46. 10.1016/j.stem.2011.12.004.

Park, I.K., Qian, D., Kiel, M., Becker, M.W., Pihalja, M., Weissman, I.L., Morrison, S.J., and Clarke, M.F. (2003). Bmi-1 is required for maintenance of adult self-renewing haematopoietic stem cells. Nature 423, 302–305. 10.1038/nature01587.

Parreno, V., Martinez, A.M., and Cavalli, G. (2022). Mechanisms of Polycomb group protein function in cancer. Cell Res 32, 231–253. 10.1038/s41422-021-00606-6.

Piunti, A., Rossi, A., Cerutti, A., Albert, M., Jammula, S., Scelfo, A., Cedrone, L., Fragola, G., Olsson, L., Koseki, H., et al. (2014). Polycomb proteins control proliferation and transformation independently of cell cycle checkpoints by regulating DNA replication. Nat Commun 5, 3649. 10.1038/ncomms4649.

Plath, K., Fang, J., Mlynarczyk-Evans, S.K., Cao, R., Worringer, K.A., Wang, H., de la Cruz, C.C., Otte, A.P., Panning, B., and Zhang, Y. (2003). Role of histone H3 lysine 27 methylation in X inactivation. Science 300, 131–135. 10.1126/science.1084274.

Plys, A.J., Davis, C.P., Kim, J., Rizki, G., Keenen, M.M., Marr, S.K., and Kingston, R.E. (2019). Phase separation of Polycomb-repressive complex 1 is governed by a charged disordered region of CBX2. Genes Dev 33, 799–813. 10.1101/gad.326488.119.

Ramirez, F., Ryan, D.P., Gruning, B., Bhardwaj, V., Kilpert, F., Richter, A.S., Heyne, S., Dundar, F., and Manke, T. (2016). deepTools2: a next generation web server for deep-sequencing data analysis. Nucleic Acids Res 44, W160–165. 10.1093/nar/gkw257.

Satijn, D.P., Gunster, M.J., van der Vlag, J., Hamer, K.M., Schul, W., Alkema, M.J., Saurin, A.J., Freemont, P.S., van Driel, R., and Otte, A.P. (1997). RING1 is associated with the polycomb group protein complex and acts as a transcriptional repressor. Mol Cell Biol 17, 4105–4113. 10.1128/mcb.17.7.4105.

Saurin, A.J., Shiels, C., Williamson, J., Satijn, D.P., Otte, A.P., Sheer, D., and Freemont, P.S. (1998). The human polycomb group complex associates with pericentromeric heterochromatin to form a novel nuclear domain. J Cell Biol 142, 887–898. 10.1083/jcb.142.4.887.

Schindelin, J., Arganda-Carreras, I., Frise, E., Kaynig, V., Longair, M., Pietzsch, T., Preibisch, S., Rueden, C., Saalfeld, S., Schmid, B., et al. (2012). Fiji: an open-source platform for biological-image analysis. Nat Methods 9, 676–682. 10.1038/nmeth.2019.

Schmidt, J.A., Oatley, J.M., and Brinster, R.L. (2009). Female mice delay reproductive aging in males. Biol Reprod 80, 1009–1014. 10.1095/biolreprod.108.073619.

Seif, E., Kang, J.J., Sasseville, C., Senkovich, O., Kaltashov, A., Boulier, E.L., Kapur, I., Kim, C.A., and Francis, N.J. (2020). Phase separation by the polyhomeotic sterile alpha motif compartmentalizes Polycomb Group proteins and enhances their activity. Nat Commun 11, 5609. 10.1038/s41467-020-19435-z.

Shao, Z., Raible, F., Mollaaghababa, R., Guyon, J.R., Wu, C.T., Bender, W., and Kingston, R.E. (1999). Stabilization of chromatin structure by PRC1, a Polycomb complex. Cell 98, 37–46. 10.1016/S0092-8674(00)80604-2.

Shen, Y., Yue, F., McCleary, D.F., Ye, Z., Edsall, L., Kuan, S., Wagner, U., Dixon, J., Lee, L., Lobanenkov, V.V., and Ren, B. (2012). A map of the cis-regulatory sequences in the mouse genome. Nature 488, 116–120. 10.1038/nature11243.

Shinohara, T., Orwig, K.E., Avarbock, M.R., and Brinster, R.L. (2000). Spermatogonial stem cell enrichment by multiparameter selection of mouse testis cells. Proc Natl Acad Sci U S A 97, 8346–8351. 10.1073/pnas.97.15.8346.

Silva, J., Mak, W., Zvetkova, I., Appanah, R., Nesterova, T.B., Webster, Z., Peters, A.H., Jenuwein, T., Otte, A.P., and Brockdorff, N. (2003). Establishment of histone h3 methylation on the inactive X chromosome requires transient recruitment of Eed-Enx1 polycomb group complexes. Dev Cell 4, 481–495. 10.1016/s1534-5807(03)00068-6.

Skarnes, W.C., Rosen, B., West, A.P., Koutsourakis, M., Bushell, W., Iyer, V., Mujica, A.O., Thomas, M., Harrow, J., Cox, T., et al. (2011). A conditional knockout resource for the genome-wide study of mouse gene function. Nature 474, 337–342. 10.1038/nature10163.

Strom, A.R., Emelyanov, A.V., Mir, M., Fyodorov, D.V., Darzacq, X., and Karpen, G.H. (2017). Phase separation drives heterochromatin domain formation. Nature 547, 241–245. 10.1038/nature22989.

Struhl, G., and Akam, M. (1985). Altered distributions of Ultrabithorax transcripts in extra sex combs mutant embryos of Drosophila. EMBO J 4, 3259–3264.

Stuart, T., Butler, A., Hoffman, P., Hafemeister, C., Papalexi, E., Mauck, W.M., 3rd, Hao, Y., Stoeckius, M., Smibert, P., and Satija, R. (2019). Comprehensive Integration of Single-Cell Data. Cell 177, 1888–1902 e1821. 10.1016/j.cell.2019.05.031.

Takada, Y., Isono, K., Shinga, J., Turner, J.M., Kitamura, H., Ohara, O., Watanabe, G., Singh, P.B., Kamijo, T., Jenuwein, T., et al. (2007). Mammalian Polycomb Scmh1 mediates exclusion of Polycomb complexes from the XY body in the pachytene spermatocytes. Development 134, 579–590. 10.1242/dev.02747.

Tatavosian, R., Kent, S., Brown, K., Yao, T., Duc, H.N., Huynh, T.N., Zhen, C.Y., Ma, B., Wang, H., and Ren, X. (2019). Nuclear condensates of the Polycomb protein chromobox 2 (CBX2) assemble through phase separation. J Biol Chem 294, 1451–1463. 10.1074/jbc.RA118.006620.

Wang, H., Wang, L., Erdjument-Bromage, H., Vidal, M., Tempst, P., Jones, R.S., and Zhang, Y. (2004). Role of histone H2A ubiquitination in Polycomb silencing. Nature 431, 873–878. 10.1038/nature02985.

Wang, J., Mager, J., Chen, Y., Schneider, E., Cross, J.C., Nagy, A., and Magnuson, T. (2001). Imprinted X inactivation maintained by a mouse Polycomb group gene. Nat Genet 28, 371–375. 10.1038/ng574.

Wei, C., Lin, H., and Cui, S. (2018). The Forkhead Transcription Factor FOXC2 Is Required for Maintaining Murine Spermatogonial Stem Cells. Stem Cells Dev 27, 624–636. 10.1089/scd.2017.0233.

Xie, H., Xu, J., Hsu, J.H., Nguyen, M., Fujiwara, Y., Peng, C., and Orkin, S.H. (2014). Polycomb repressive complex 2 regulates normal hematopoietic stem cell function in a developmental-stage-specific manner. Cell Stem Cell 14, 68–80. 10.1016/j.stem.2013.10.001.

Yoshida, S., Sukeno, M., and Nabeshima, Y. (2007). A vasculature-associated niche for undifferentiated spermatogonia in the mouse testis. Science 317, 1722–1726. 10.1126/science.1144885.

Zhang, Y., Liu, T., Meyer, C.A., Eeckhoute, J., Johnson, D.S., Bernstein, B.E., Nusbaum, C., Myers, R.M., Brown, M., Li, W., and Liu, X.S. (2008). Model-based analysis of ChIP-Seq (MACS). Genome Biol 9, R137. 10.1186/gb-2008-9-9-r137.

Zhou, Q., Li, Y., Nie, R., Friel, P., Mitchell, D., Evanoff, R.M., Pouchnik, D., Banasik, B., McCarrey, J.R., Small, C., and Griswold, M.D. (2008). Expression of stimulated by retinoic acid gene 8 (Stra8) and maturation of murine gonocytes and spermatogonia induced by retinoic acid in vitro. Biol Reprod 78, 537–545. 10.1095/biolreprod.107.064337.

