## Supplementary material for "Cell type-specific role of CBX2-cPRC1 at the onset of spermatogonial differentiation": Key resources table

| REAGENT or RESOURCE | SOURCE | IDENTIFIER |
|---|---|---|
| Antibodies | | |
| Rabbit monoclonal anti **HA-tag** (clone C29F4)<br>- IF paraffin section 1:300<br>- IF wholemount tubule 1:1,000<br>- Western 1:10,000<br>- Cut&Run 1:1,000 | Cell Signaling Technology | RRID:AB_1549585;<br>Cat# 3724 |
| Mouse monoclonal anti **HA-tag** (clone 16B12)<br>- IF dissociated cells on coverslips 1:200<br>- Western 1:1,000<br>- Cut&Run 1:50 | Biolegend | RRID:AB_2820200<br>Cat#: 901516 |
| Rabbit polyclonal anti **HA-tag**<br>- Cut&Run 1:200 | Abcam | RRID:AB_307019<br>Cat #: ab9110 |
| Mouse monoclonal anti **SALL4** (clone EE-30)<br>- IF paraffin section 1:200<br>- IF wholemount tubule 1:600 | Santa Cruz Biotechnology | RRID:AB_1129262;<br>Cat# sc-101147 |
| Sheep polyclonal anti **FOXC2**<br>- IF wholemount tubule 1:1,200 | R&D Systems | RRID:AB_10973139<br>Cat# AF6989 |
| Rabbit monoclonal anti **CCND2** (clone D52F9)<br>- IF wholemount tubule 1:400 | Cell Signaling Technology | RRID:AB_2070685<br>Cat# 3741 |
| Rabbit monoclonal anti **Cleaved Caspase 3 (Asp175)** (clone D3E9)<br>- IF wholemount tubule 1:400 | Cell Signaling Technology | RRID:AB_10897512<br>Cat# 9579 |
| Rabbit polyclonal anti **CBX2** (rabbit 855, affinity purified)<br>- IF dissociated cells on coverslips 1:100<br>- Cut&Run: day91 serum 1:1,000 | This paper | N/A |
| Rabbit polyclonal anti **CBX2** (rabbit 856, affinity purified)<br>- IF dissociated cells on coverslips 1:100<br>- Cut&Run: day91 serum 1:1,000 | This paper | N/A |
| Rabbit polyclonal anti **CBX4**<br>- Western: 1:2,000 | Abcam | RRID: N/A<br>Cat# ab139815 |
| Rabbit polyclonal anti **CBX8**<br>- Western: 1:10,000 | Bethyl Laboratories | RRID:AB_2071525<br>Cat# A300-882A |
| Rabbit polyclonal anti **RING1B**<br>- Western: 1:50,000 | Bethyl Laboratories | RRID:AB_10632773<br>Cat# A302-869A |
| Rabbit monoclonal anti **RING1B** (clone D22F2)<br>- IF dissociated cells on coverslips 1:1,000<br>- Cut&Run: 1:500 | Cell Signaling Technology | RRID: N/A<br>Cat# 5694 |
| Mouse monoclonal **PHC2**<br>- IF dissociated cells on coverslips 1:100 | Active Motif | RRID:AB_2615060<br>Cat# 39661 |
| Rabbit **PHC2**<br>- Cut&Run: 1:200 | Cell Signaling Technology | RRID: N/A<br>Cat# H8523 |
| Rabbit polyclonal anti **BMI1**<br>- Cut&Run: 1:200 | Bethyl Laboratories | RRID:AB_1210891<br>Cat# A301-694A |
| Rabbit monoclonal anti **H3K27me3** (clone C36B11)<br>- Cut&Run: 1:500 | Cell Signaling Technology | RRID:AB_2616029<br>Cat# 9733 |
| Rabbit polyclonal anti **H3K4me3**<br>- Cut&Run: 1:1,000 | Millipore | RRID:AB_1977252<br>Cat# 07-473 |
| Rat monoclonal anti-c-Kit (CD117), conjugated with PE (clone 3C11)<br>- Flow Cytometry: 1:1,000 | Miltenyi Biotec | RRID:AB_2801976<br>Cat#: 130-122-937 |

| | | |
|---|---|---|
| Donkey polyclonal anti **Mouse**, conjugated with Alexa Fluor 488<br>- IF (all types): 1:1,000 | Thermo Fisher Scientific | RRID:AB_141607<br>Cat# A21202 |
| Donkey polyclonal anti **Mouse**, conjugated with Alexa Fluor 568<br>- IF (all types): 1:1,000 | Thermo Fisher Scientific | RRID:AB_2534013<br>Cat# A10037 |
| Donkey polyclonal anti **Rabbit**, conjugated with Alexa Fluor 488<br>- IF (all types): 1:1,000 | Thermo Fisher Scientific | RRID:AB_253579<br>Cat# A21206 |
| Donkey polyclonal anti **Rabbit**, conjugated with Alexa Fluor 568<br>- IF (all types): 1:1,000 | Thermo Fisher Scientific | RRID:AB_2534017<br>Cat# A10042 |
| Donkey polyclonal anti **Sheep**, conjugated with Alexa Fluor 488<br>- IF (all types): 1:500 | Jackson ImmunoResearch | RRID:AB_2340745<br>Cat# 713-545-147 |
| Donkey polyclonal anti **Sheep**, conjugated with Alexa Fluor 647<br>- IF (all types): 1:500 | Jackson ImmunoResearch | RRID:AB_2340751<br>Cat# 713-605-147 |
| Donkey polyclonal anti **Mouse**, conjugated with HRP<br>- Western: 1:10,000 | Jackson ImmunoResearch | RRID:AB_2340770<br>Cat# 715-035-150 |
| Donkey polyclonal anti **Rabbit**, conjugated with HRP<br>- Western: 1:10,000 | Jackson ImmunoResearch | RRID:AB_10015282<br>Cat# 711-035-152 |
| Bacterial and virus strains | | |
| Biological samples | | |
| Chemicals, peptides, and recombinant proteins | | |
| Tamoxifen | Sigma-Aldrich | Cat# T5648 |
| Critical commercial assays | | |
| Chromium Single Cell 3’ Library & Gel Bead Kit v2 | 10X Genomics | Cat# 120267 |
| Chromium Single Cell A Chip Kit | 10X Genomics | Cat# 1000009 |
| Chromium Next GEM Single Cell 5' Kit v2 | 10X Genomics | Cat# 1000265 |
| Chromium Next GEM Chip K Single Cell Kit | 10X Genomics | Cat# 1000287 |
| Deposited data | | |
| Raw and processed high throughput sequencing data (Cut&Run, scRNA-seq) | This paper | NCBI GEO accession: GSE210369 |

| | | |
|---|---|---|
| Experimental models: Cell lines | | |
| Experimental models: Organisms/strains | | |
| Mouse: C57BL/6J | The Jackson Laboratory | RRID:IMSR_JAX:000664; Cat# 000664 |
| Mouse: Cbx2$^{2xHA}$ | This paper | N/A |
| Mouse: Cbx2$^{\Delta}$ | This paper | N/A |
| Mouse: Cbx2$^{23KRA}$ | Lau *et al.*, 2017 Maintained in lab colony Available through MMRRC | RRID:MMRRC_050534-UNC Cat# 050534-UNC |
| Mouse: Cbx2$^{Flox}$ (Cbx2tm1a(KOMP)Wtsi) | UC Davis Mouse Biology Program Knock Out Mouse Project (KOMP) | RRID:MMRRC_046927-UCD |
| Mouse: CAGG's Flpo (C57BL/6N-Tg(CAG-Flpo)1Afst/Mmucd) | UC Davis Mutant Mouse Resource Research Center (MMRRC) | RRID:MMRRC_036512-UCD |
| Mouse: R26-CreERT2 (B6.129-Gt(ROSA)26Sortm1(cre/ERT2)Tyj/J) | The Jackson Laboratory | RRID:IMSR_JAX:008463 Cat# 008463 |
| Oligonucleotides | | |
| Recombinant DNA | | |
| Software and algorithms | | |
| Fiji | Schindelin *et al.*, 2012 | https://imagej.net/software/fiji/ |
| Matlab | Mathworks | https://www.mathworks.com/products/matlab.html |
| R | The R project | https://www.r-project.org/ |
| Trim Galore | Martin *et al.*, 2012 | https://www.bioinformatics.babraham.ac.uk/projects/trim_galore/ |

| | | |
|---|---|---|
| Bowtie2 | Langmead and Salzberg, 2012 | http://bowtie-bio.sourceforge.net/bowtie2/index.shtml |
| Samtools | Li *et al.*, 2009 | http://www.htslib.org/ |
| Picard | Broad Institute | http://broadinstitute.github.io/picard |
| Deeptools | Ramirez *et al.*, 2016 | https://deeptools.readthedocs.io/en/develop/ |
| MACS2 | Zhang *et al.*, 2008 | https://github.com/macs3-project/MACS |
| Cell ranger | 10X Genomics | https://support.10xgenomics.com/single-cell-gene-expression/software/pipelines/latest/installation |
| Seurat | Stuart *et al.*, 2019; Hao *et al.*, 2021 | https://satijalab.org/seurat/ |
| All other scripts | This paper | https://github.com/jongminkmg/Cbx2 |
| Other | | |
